# NKG2C^+^CD27^−^ Defines Human CD8^+^ Regulatory T Cells

**DOI:** 10.64898/2026.06.15.732401

**Authors:** Xuyang Li, Somen K. Mistri, Tian Cao, Rajesh K. Krishnan, Martin Hemberg, Howard L. Weiner, Dan Hu

## Abstract

Here we identify NKG2C^+^CD27^−^ as a novel surface marker unifying multiple previously reported CD8^+^ regulatory T cell (Treg) populations. This population displays high clonality and divides into CD226^−^ (Treg1) and CD226^+^ (Treg2) subsets, with Treg2 exhibiting stronger suppressive activities. Up to 35% of CD8^+^ T_EMRA_ cells are Tregs, whereas approximately 85% of CD8^+^ Tregs are T_EMRA_ cells, which increase with aging. These findings establish a unified and novel cell surface marker for CD8^+^ Tregs and their subsets, which resolves prior heterogeneity in the field, and show that CD8^+^ T_EMRA_ cells are a heterogeneous population that includes T cells with regulatory function. Our findings provide a critical framework for the isolation and in-depth functional characterization of CD8^+^ Tregs in health, aging, and disease.

## INTRODUCTION

Regulatory T cells (Tregs) are a specialized subset of T cells known for their immunosuppressive properties, and they are essential for maintaining immune homeostasis and self-tolerance. The concept of suppressor T cells originated in 1970 when Gershon and Kondo discovered that certain T cells could dampen immune responses (*1*). These were called “suppressor” T cells and were extensively studied in the 1970s and 1980s as a proposed mechanism of immune regulation, with most of the suppressor T cells identified as CD8^+^, though some CD4^+^ cells also exhibited suppressive functions (*2*). The I-J molecule was thought to be crucial for CD8^+^ suppressor T cell function, but its genetic basis was disproven in the mid-1980s, leading to a decline in suppressor T cell research (*3*). Furthermore, early studies were also hampered by poor reproducibility and lack of clear cell surface markers, making it difficult to investigate suppressor T cells as a distinct lineage (*4*). The suppressor T cell concept was then better defined with the identification of CD25^+^CD4^+^ regulatory T cells (Tregs) in 1995 by Sakaguchi and his colleagues (*5*), which could be reliably distinguished and shown to play a crucial role in immune tolerance and homeostasis. A subsequent landmark achievement in CD4⁺ Treg studies was the discovery of Foxp3 as their defining transcription factor in 2000s (*6, 7*). These breakthroughs have established CD4^+^ Tregs as dominant players—not only in immunosuppressive T cell research but also in immunology and immune-related diseases.

In contrast, regulatory CD8^+^ T cell studies have not attracted widespread attention despite data accumulated in past decades which have supported the existence of CD8^+^ Tregs (*8, 9*). A significant breakthrough in CD8^+^ Treg research was the discovery of murine Qa-1-restricted CD8^+^ Tregs (*10–13*), which expressed Ly49 (*14*), a family of C-type lectin-like receptors interacting with MHC class I molecules (*15*). In addition, Helios is a key regulator of CD8⁺Ly49^+^ Treg function; while not Treg-specific, it promotes transcriptional programs associated with suppressive activity and is upregulated in CD8⁺ Tregs (*16*). Humans do not have Ly49 genes; instead killer cell immunoglobulin-like receptors (KIRs) are used to perform similar function (*17*). CD8^+^KIR^+^ Tregs (*18*) and CD8^+^CD56^+^CD161^−^ Tregs (*19, 20*) are found exclusively in humans, given that neither KIRs nor CD56 are expressed by immune cells in mice. In contrast, CD8^+^CD122^+^, CD8^+^CD28^−/low^ and CD8^+^Foxp3^+^ Tregs have been reported in both species (*9, 21, 22*). Among the various types of CD8^+^ Tregs, the relationship between them is not well-defined, except for Ly49^+^ Tregs in mice and KIR^+^ Tregs in humans, which are considered mouse-human counterparts (*18*).

Given the diverse CD8^+^ Treg subsets reported, they are considered a heterogeneous population as opposed to well-defined CD4^+^ Tregs. Although CD8^+^Ly49^+^ Tregs are CD122^+^, only ∼20% of CD8^+^CD122^+^ T cells are Ly49^+^ Tregs, compared with fewer than 5% of total CD8^+^ T cells (*14*). This observation indicates that Ly49^+^ Tregs are enriched within the CD122^+^ subset, although most cells are non-Tregs. Similarly, because CD28 is expressed on naïve CD8⁺ T cells and downregulated upon activation, most human CD8⁺ T cells are CD28^−^ (*23*); accordingly, Tregs are most likely enriched in the CD28^−/low^ subset but still represent a minority. Moreover, CD8^+^CD56^+^CD161^−^ Tregs and CD8^+^KIR^+^ Tregs show similarities in cell surface markers and function (*18–20, 24–26*). Therefore, an important question is whether these diverse CD8^+^ Treg subsets represent overlapping, rather than distinct populations.

Apart from lineage diversity, T cells can also be categorized by differentiation stages: naïve, effector, memory, and terminally differentiated (*27, 28*). CD45RA is a glycoprotein, co-expressed with CD27 and CCR7 on human naïve T cells. Memory T cells are classified as CD45RA^−^CD27^−^CCR7^−^ (*29*). In humans, Tcells (T effector memory cells re-expressing CD45RA) are a subset of memory T cells that re-acquire CD45RA expression after prior activation (*27, 29, 30*). However, these cells still maintain the CD27^−^CCR7^−^ phenotype (*27, 29, 31*). CD8^+^ Tcells are commonly regarded as terminally differentiated, highly cytolytic cells that function much like pro-inflammatory cytotoxic T lymphocytes (CTLs) (*32, 33*). They are associated with human diseases including chronic infections, autoimmune disorders (*34–36*) and neurodegeneration and neuroinflammation (*37–43*). Interestingly, most previously reported CD8^+^CD56^+^CD161^−^ Tregs display a Tphenotype characterized by CD45RA^+^CD45RO^−^CD27^−^CCR7^−^ expression (*19, 20*). These results suggest a previously unappreciated heterogeneity within the CD8^+^ Tcompartment; disease-associated CD8^+^ Tcells may include both proinflammatory CTLs and immunosuppressive CD8^+^ Tregs.

Here, using single-cell multi-omics, high-dimensional flow cytometry and functional analysis, we comprehensively characterized human CD8^+^ Tregs and identified NKG2C^+^CD27^−^ as a defining surface marker signature that unifies previously reported CD8^+^ Treg populations into a single entity. CD8^+^ Tregs comprised two distinct subtypes: CD226^−^PTGDS^−/low^ (Treg1) and CD226^+^PTGDS^hi^ (Treg2), with Treg2 exhibiting stronger suppressive activity compared to Treg1.

## RESULTS

### CD8^+^CD56^+^CD161^−^ and CD8^+^KIR^+^ T cells overlap and are both enriched in CD8^+^ Tcells

We have previously postulated that, given the substantial similarities in function and cell surface marker expression between CD8^+^CD56^+^CD161^−^ regulatory T cells (*20*) and CD8^+^KIR^+^ regulatory T cells (*18*), there is an overlap between these two populations (*24, 25*). Additionally, the majority of CD8^+^CD56^+^CD161^−^ T cells (hereafter referred to as CD8^+^CD56^+^ cells) display a Tphenotype of CD45RA^+^CD45RO^−^CD27^−^CCR7^−^ (*20*). These prior observations prompted us to investigate the degree of overlap among CD8^+^CD56^+^ cells, CD8^+^KIR^+^ cells and CD8^+^ Tcells. We performed flow cytometry on PBMCs from 20 healthy donors using a 13-antibody panel that identified these three populations (Fig. 1A-C and fig. S1A), which included antibodies recognizing CD3, CD4, CD8, CD45RA, CD45RO, CD56, CD161, CCR7, CD27, KIR2DL2/L3/S2, KIR3DL1, PD-1 and CXCR6 (table S1). We defined CD8^+^KIR^+^ cells as CD8^+^ T cells positive for KIR2DL2/L3/S2 and/or KIR3DL1 (Fig. 1B), using the same antibody clones originally used to identify regulatory CD8^+^KIR^+^ cells (*18*). We simplified the Tmarker definition from CD45RA^+^CD27^−^CCR7^−^ to CD45RA^+^CD27^−^, based on the observation that CD45RA^+^CD27^−^ T cells were CCR7^−^ (Fig. 1C). Virtually all Tcells were found within the CD8^+^ T cell compartment: 0.11% of CD4^+^ T cells and 6.1% of CD8^+^ T cells were Tcells (paired, two-tailed *t*-test, *p* < 0.0001; fig. S1B). The frequencies of naïve T (Tn) cells were similar between CD4^+^ and CD8^+^ T cells (fig. S1C), whereas the frequencies of central memory T (Tcm) cells were significantly higher in CD4^+^ T cells (paired, two-tailed *t*-test, *p* < 0.0001; fig. S1D). Interestingly, although we gated on CD161^−^ cells to define CD8^+^CD56^+^ cells, these cells—like CD8^+^ Tcells and CD8^+^KIR^+^ cells—expressed higher levels of CD161 compared to Tcm cells (Ordinary one-way ANOVA, Dunnett’s multiple comparisons test, *p* < 0.0001) (fig. S1E), suggesting that most CD8^+^ T cells have a basal level of CD161 expression, which we also observed in subsequent single-cell RNA-seq (scRNA-seq) analysis (Fig. 2B). Therefore, CD8^+^CD56^+^ cells are CD161^low^ rather than CD161^−^.

**Fig. 1.**
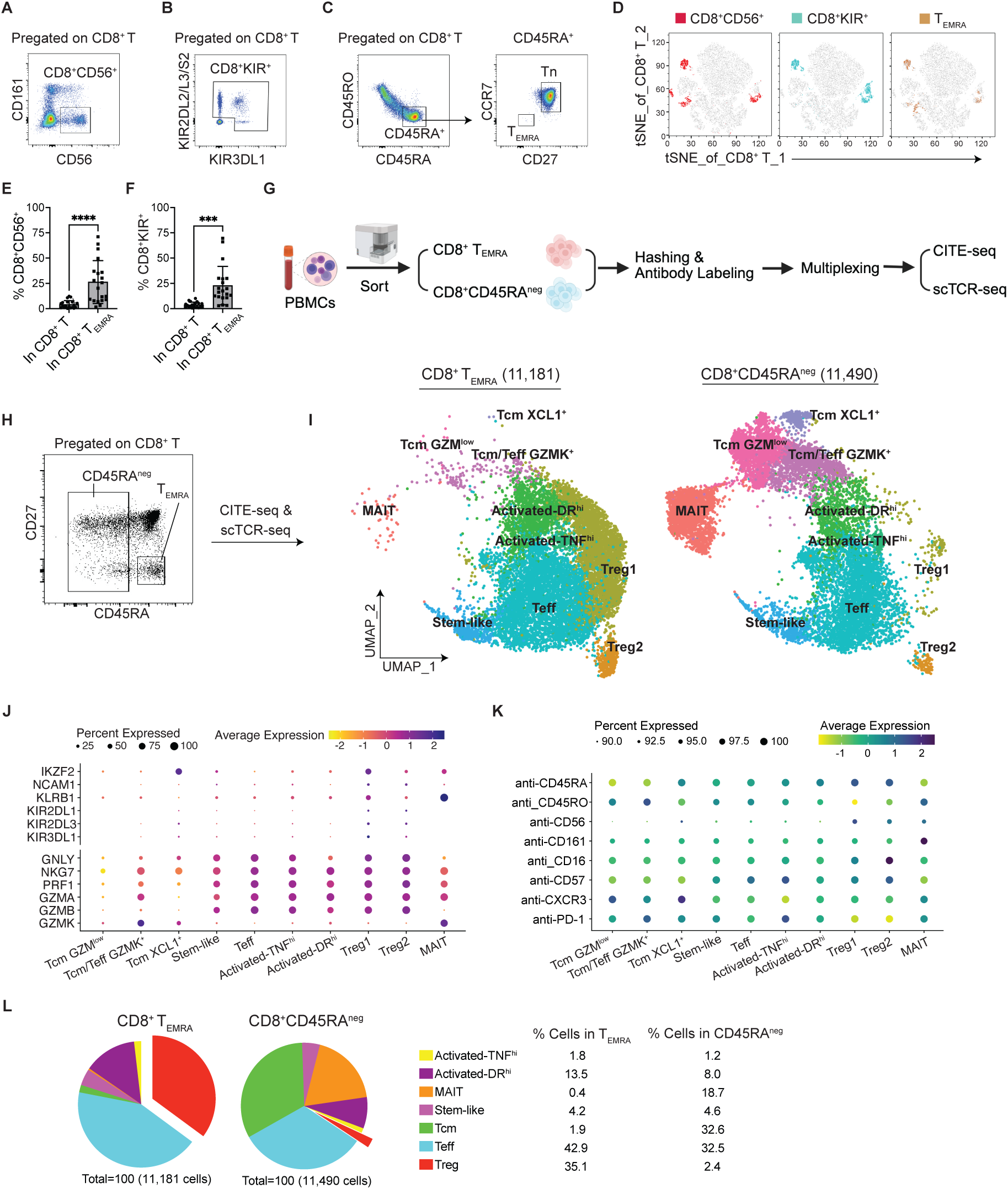
CD8^+^ Tregs are highly enriched within the Tcells population and can be divided into two distinct subsets. (**A**-**F**) PBMCs from healthy donors (HCs; n = 20) were analyzed using a CyTek® Aurora flow cytometer with a 13-antibody panel to assess the frequencies of CD56^+^CD161^−^, KIR^+^, and T subsets within the CD8^+^ T cell population. (**A-C**) Representative plots illustrating the gating strategy for CD56^+^CD161^−^ (**A**), KIR^+^ (**B**), and T(**C**) subsets within the CD8^+^ T cell compartment. **D**) tSNE plot of CD8^+^ T cells from a representative donor showing that CD8^+^ CD56^+^CD161^−^, KIR^+^ and Tsubsets represent overlapping populations. (**E, F**) Both CD8^+^CD56^+^ (**E**) and CD8^+^KIR^+^ cells (**F**) were highly enriched within the CD8^+^ Tcompartment (*n* = 20; mean ± s.d.). Paired, two-tailed, *t*-test. \*\*\**p* = 0.0001, **** *p* < 0.0001. (**G**-**L**) CITE-seq and scTCR-seq of CD8^+^ Tand CD45RA^neg^ cells isolated from PBMCs of HCs (*n* = 4; 2 females, 2 males). (**G**) Scheme of experimental design: mix-sample strategy to enrich CD8^+^ Tcells by combining equal numbers of CD8^+^ Tand CD45RA^neg^ cells from each donor for CITE-seq and scTCR-seq. (**H**) Representative plot showing gating on CD8^+^ T cells for CD8^+^ Tand CD45RA^neg^ cell sorting. (**I**) UMAP plots split by isolated CD8^+^ T(*n* = 11,181) and CD45RA^neg^ (*n* = 11,490) cells, colored by the 10 identified clusters. (**J**) Expression of CD8^+^ Treg-associated and cytotoxicity-related genes across CD8^+^ T cell clusters. Dot color indicates the level of gene expression and dot size indicates the proportion of cells expressing the gene. (**K**) Cell surface protein expression in CD8^+^ T cell clusters measured by ADT. Dot color indicates the level of gene expression and dot size indicates the percentage of cells expressing the gene. (**L**) Cell frequency of CD8^+^ T cell clusters in sorted CD8^+^ Tand CD45RA^neg^ cells, respectively. Tcm represents the combined Tcm GZM^low^, Tcm/Teff GZMK^+^ and Tcm XCL1^+^ populations. Treg represents the combined Treg1 and Treg2 populations.

**Fig. 2.**
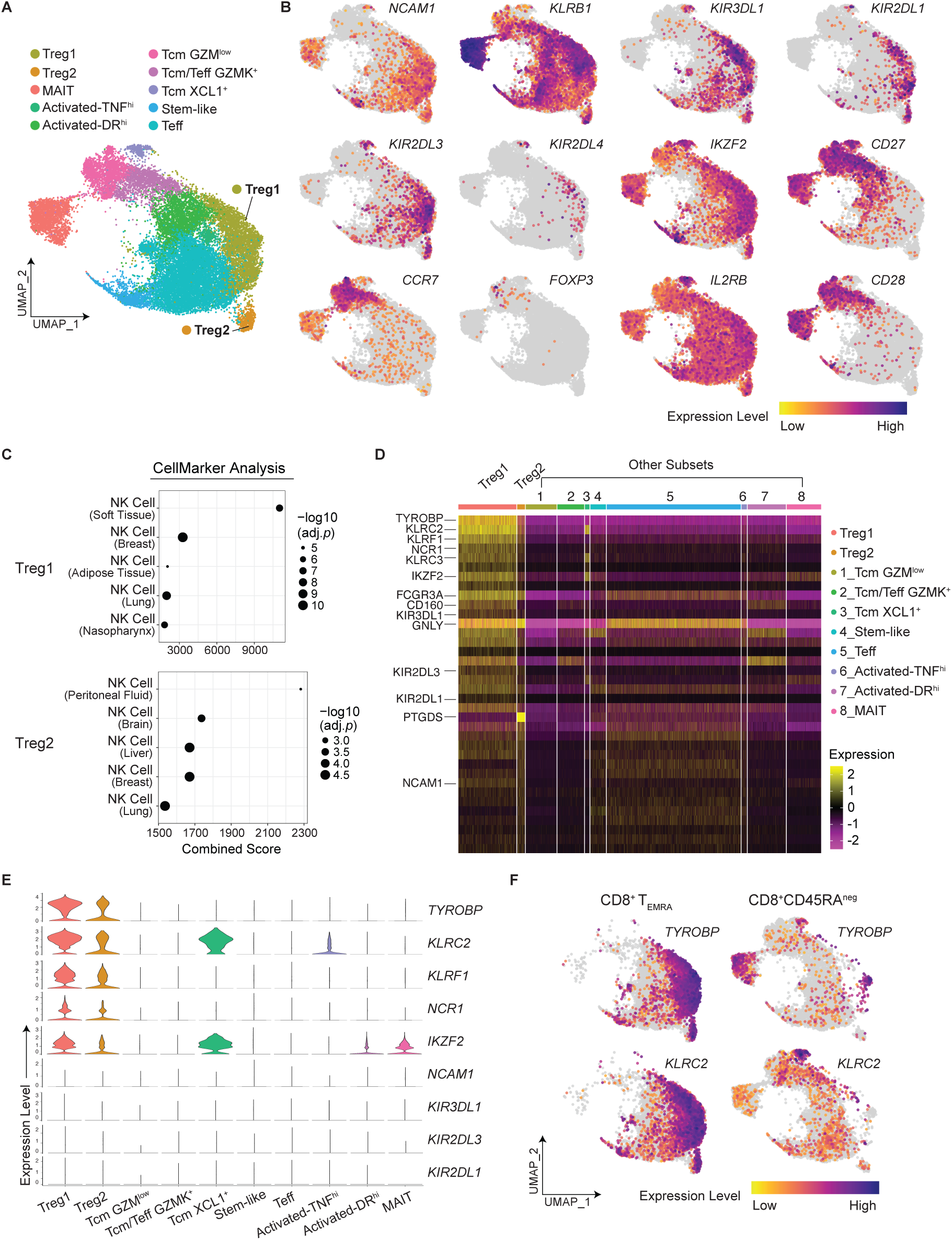
Identification of TYROBP and NKG2C (*KLRC2*) as novel and improved markers for CD8^+^ Tregs. CITE-seq of CD8^+^ Tand CD45RA^neg^ cells (continued). (**A**) UMAP plot of merged CD8^+^ Tand CD45RA^neg^ cells showing the distribution of CD8^+^ T cell subsets. (**B**) UMAPs showing the expression of previously reported CD8^+^ Treg marker genes and other lineage-defining genes. (**C**) The top five hits identified by the CellMarker Augmented 2024 analysis in Enrichr for Treg1 (Upper panel) and Treg2 cells (Lower panel). (**D**) Heat map showing expression of the top 20 up-regulated genes in CD8^+^ Treg1 and Treg2 cells across CD8^+^ T cell subsets. Genes were ranked based on a composite score calculated as the fold change multiplied by the percentage of positive cells. (**E**) Violin plots comparing the expression of the top 4 up-regulated genes in CD8^+^ Treg1 with previously reported CD8^+^ Treg-marker genes (*IKZF2*, *NCAM1* and *KIR* genes). (**F**) UMAPs showing the expression of *TYROBP* and *KLRC2* in CD8^+^ Tand CD45RA^neg^ cells, respectively.

We then performed t-distributed stochastic neighbor embedding (t-SNE) analysis on CD8^+^ T cells and visualized the distribution of CD8^+^CD56^+^, CD8^+^KIR^+^ and CD8^+^ Tcells. The t-SNE plot revealed substantial overlap among the three populations (Fig. 1D). On average, 4.1% of total CD8^+^ T cells were CD56^+^CD161^−^, whereas within the CD8^+^ Tcompartment, 26.3% cells were CD56^+^CD161^−^ (paired, two-tailed *t*-test, *p* < 0.0001) (Fig. 1E). Similarly, 3.7% of total CD8^+^ T cells were KIR^+^, compared to 22.8% of CD8^+^ Tcells (paired, two-tailed *t*-test, *p* = 0.0001) (Fig. 1E). Thus, both CD8^+^CD56^+^ cells and CD8^+^KIR^+^ cells were enriched in the CD8^+^ Tcompartment relative to the total CD8^+^ T cell population. Similar pairwise enrichment was also observed in CD8^+^CD56^+^ cells (fig. S1F) as well as in CD8^+^KIR^+^ cells (fig. S1G). CD45RA expression in CD8^+^CD56^+^ cells and CD8^+^KIR^+^ cells ranged from negative to high expression levels, with a small CD45RA^−^CD45RO^+^ subset. Both populations lacked CCR7 expression, placing them outside the conventional CD45RA^+^CD27^+^CCR7^+^ naïve T cell compartment (fig. S1H).

In summary, our flow cytometric analysis revealed that CD8^+^CD56^+^, CD8^+^KIR^+^ and CD8^+^ Tcells exhibit substantial overlap, indicating they are not discrete subsets but rather share phenotypic features. Both CD8^+^CD56^+^ cells and CD8^+^KIR^+^ cells lack the key features of naïve T cells and thus fall outside the naïve T cell compartment.

### CD56 and KIRs identify identical cell clusters in single-cell multi-omics analysis

To characterize CD8^+^ Tregs transcriptome-wide, we performed CITE-seq and scTCR-seq using a mixed-sample strategy. Briefly, we isolated CD3^+^CD8^+^CD45RA^+^CD27^−^ (T) and CD3^+^CD8^+^CD45RA^−^ (CD45RA^neg^) cells from the PBMCs of four healthy adults by cell sorting and subjected an equal number of Tand CD45RA^neg^ cells from each individual to CITE-seq and scTCR-seq (Methods and Fig. 1G and H). The antibody-derived tag (ADT) panel measured the protein expression of 10 cell surface markers, including CD4, CD8, CD45RA, CD45RO, CD56, CD161, CD16, CD57, CXCR3 and PD-1 (table S2). After quality control followed by normalization and integration (Methods and fig. S2A-D), high-quality singlets (11,181 for Tcells and 11,490 for CD45RA^neg^ cells) extracted from the CITE-seq dataset were clustered and visualized with uniform manifold approximation projection (UMAP). We identified 10 CD8^+^ T cell subpopulations in our dataset (Fig. 1I), based on CD8^+^ Treg marker gene and TCR gene expression, and protein expression detected by the antibody-derived tags (ADTs) (Fig. 1J and Fig. 1K, fig. S2E and S2F). We found that, CD8^+^ Treg1 and Treg2 clusters were identified by high RNA expression of *NCAM1* (encodes CD56), *KIR2DL1*, *KIR2DL3* and *KIR3DL1* (Fig. 1J) and by high surface protein expression of CD45RA and CD56 and low surface protein expression of CD161 detected by ADTs (Fig. 1K). Of note, *KIR2DL1*, *KIR2DL3* and *KIR3DL1* were the three most highly expressed KIR genes in our dataset. Additionally, although Helios is induced during T cell activation and proliferation, it is not a Treg-specific marker (*44*), given that it is expressed in mouse CD8^+^ Tregs (*16*) and human CD8^+^KIR^+^ cells (*18*); We used co-expression of *IKZF2* (encodes Helios) and KIR genes as one of the criteria to identify CD8^+^ Treg clusters. The co-expression of *NCAM1* (CD56), *KIR2DL1*, *KIR2DL3*, *KIR3DL1* and *IKZF2* in the Treg 1 and Treg 2 cell clusters that expressed low levels of *KLRB1* (CD161) suggests that CD8^+^CD56^+^ and CD8^+^KIR^+^ cells represent the same Treg populations.

Based on the marker genes of individual clusters, we identified central memory T (Tcm) cell clusters by high gene expression of *SELL*, *IL7R*, *TCF7* and *LEF1* and by surface protein expression of CD45RO, and subsequently divided them into Tcm GZM^low^, Tcm/Teff GZMK^+^ and Tcm XCL1^+^ based on their differential expression of various granzyme genes and *XCL1* (Fig. 1J and Fig. 1K and fig. S2E). Mucosal-associated invariant T (MAIT) cells were identified by high RNA levels of *KLRB1* (encodes CD161) (Fig. 1J), by expression of *TRAV1-2* and *TRAJ33/20/12* (encode the semi-invariant α chains of MAIT cells) (fig. S2F) and by high surface protein expression of CD161 (Fig. 1K). Remaining cell clusters, including the stem-like, Teff (T effector), activated-TNF^hi^ and activated-(HLA-)DR^hi^ clusters, were assigned by integrated assessment of T-cell activation markers (CD45RO, HLA-DR, CD57, granzymes, cytokines, etc.) and pathway enrichment analysis of cluster markers using the bioinformatics platform Enrichr (*45–47*). The high resolution of CITE-seq and enhanced CD8^+^ Treg signals by mixing equal numbers of CD8^+^ Tand CD45RA^neg^ cells (the mix sample strategy) allowed us to identify two distinct CD8^+^ Treg populations: Treg1 and Treg2 (Fig. 1I). Consistent with our flow cytometry results, CD8^+^ Tregs were highly enriched in the CD8^+^ Tcompartment (35.1% of Tcells), while a subset of CD8^+^ Tregs was also found among CD45RA^neg^ cells (2.4% of CD45RA^neg^ cells) (Fig. 1L). Tcm cells and MAIT cells were primarily observed in the CD45RA^neg^ compartment. In contrast to the biased distribution of Tregs, Tcm and MAIT cells, Teff, stem-like and activated T cells showed a balanced distribution between CD8^+^ Tand CD45RA^neg^ populations (Fig. 1L), all expressing high levels of cytolytic (cytotoxic) effector genes *GZMA*, *GZMB*, *PRF1* and *NKG7* (Fig. 1J).

Taken together, by increasing the representation of CD8^+^ Tregs through equal mixing CD8^+^ Tand CD8^+^CD45RA^neg^ cells in the CITE-seq analysis, we found that CD56 and KIRs identified the same cell clusters. This indicates that CD8^+^CD56^+^ and CD8^+^KIR^+^ cells represent the same Treg population, which can further be subdivided into two transcriptionally distinct subsets. Consistent with our flow cytometric analysis above, CD8^+^ Tregs are highly enriched within the CD8^+^ Tcompartment.

### CD8^+^ Treg1 and Treg2 clusters unify prior CD8^+^ Treg phenotypes

In addition to the previously reported human CD8^+^ Treg phenotypes—namely CD8^+^CD56^+^ and CD8^+^KIR^+^—other populations such as CD8^+^CD122^+^(*48*) and CD8^+^CD28^−/low^ (*49*) have also been reported. Notably, both CD8^+^CD56^+^ and CD8^+^KIR^+^ cells exhibit regulatory activities that are independent of Foxp3 (*19, 26*), whereas CD8^+^CD56^+^ cells express low or absent levels of cell surface CD27, CD28 and CCR7 (*20*). To further explore the overlap among these previously reported CD8^+^ Treg populations, we analyzed our CITE-seq data by first visualizing the distribution of various CD8^+^ T cell subsets within the merged CD8^+^ Tand CD45RA^neg^ populations with UMAP (Fig. 2A). We then generated feature plots for genes that have been reported as marker genes for CD8^+^ Tregs, including *NCAM1* (encodes CD56), *KIR3DL1*, *KIR2DL1*, *KIR2DL3*, *KIR2DL4*, *IKZF2* (encodes Helios) and *IL2RB* (encodes CD122), and for those known to be absent or minimally expressed in these cells, including *KLRB1* (encodes CD161), *CD27*, *CCR7*, *FOXP3* and *CD28* (Fig. 2B). These feature plots demonstrated that the Treg1 and Treg2 clusters exhibited the phenotypes of CD8^+^CD56^+^ cells (*NCAM1^+^KLRB1^low^CD27*^−^*CD28*^−^*CCR7*^−^), CD8^+^KIR^+^ cells (*KIR^+^IKZF2^+^*), CD8^+^CD122^+^ cells (*IL2RB*^+^) and CD8^+^CD28^−/low^ (*CD28*^−^), suggesting that these previously reported CD8^+^ Treg populations likely belong to the same functional subset. Nevertheless, the feature plots revealed that non-Treg clusters also expressed the Treg-associated genes *NCAM1*, *KIRs*, *IKZF2* and *IL2RB*. Notably, *IKZF2* and *IL2RB* showed broadly distributed expression among non-Treg clusters. Moreover, in line with previous reports that genetic, epigenetic, and environmental factors collectively determine the number, expression patterns, and levels of KIR genes (*17*), we also found that KIR expression in Treg clusters varied substantially across individuals (fig. S3A). Taken together, these observations suggest that the Treg1 and Treg2 clusters consolidate multiple previously reported CD8^+^ Treg populations, including CD8^+^CD56^+^, CD8^+^KIR^+^, CD8^+^CD122^+^ and CD8^+^CD28^−/low^ Tregs. Nonetheless, more Treg-specific and consistent markers are needed to define these functional T cell populations, which we describe below.

### Identification of TYROBP and NKG2C (*KLRC2*) as novel markers for CD8^+^ Tregs

To identify a specific molecular marker for CD8^+^ Tregs, we examined the marker genes of Treg1 and Treg2 clusters. We submitted their marker genes to the bioinformatics platform Enrichr (*45–47*) to conduct cell marker analysis. The top five hits identified by the CellMarker Augmented 2024 analysis in Enrichr revealed that both Treg1 and Treg2 cells closely resemble NK cells in surface marker expression, with highly significant adjusted *p*-values (*adj.p*) (Fig. 2C). A heat map was used to visualize expression of the top 20 upregulated marker genes for Treg1 and Treg2 cells across CD8^+^ T cell subsets (Fig. 2D). *TYROBP* was the top upregulated marker for Treg1, whereas *PTGDS* was the most highly upregulated marker for Treg2. These upregulated genes included a substantial number of NK-associated genes. In CD8^+^ Treg1 cells the top-ranked NK-associated genes, from high to low, were *TYROBP*, *KLRC2*, *KLRF1*, *NCR1*, *KLRC3*, *FCGR3A*, *CD160*, *KIR3DL1*, *GNLY*, *KIR2DL3*, *KIR2DL1* and *NCAM1* (Fig. 2D). Surprisingly, the previously reported CD8^+^ Treg-marker genes *KIR*s and *NCAM1* ranked near or at the bottom of the list.

To further assess the specificity of the novel CD8^+^ Treg-specific markers we identified, we generated violin plots for the top 4 marker genes of CD8^+^ Treg1 as well as previously reported CD8^+^ Treg-marker genes *IKZF2, NCAM1* and the three highest-expressed *KIR* genes. The expression of *NCAM1* and *KIR* genes was detected in less than 25% of cells in the CD8^+^ Treg1 and Treg2 clusters; by contrast, the expression of *TYROBP*, *KLRC2*, *KLRF1* and *IKZF2* was detected in more than 70% of Tregs (Fig. 2E). In contrast to the *KIR* genes, the expression of *TYROBP* and *KLRC2* was consistent across individuals (fig. S3B). TYROBP, also known as DAP12, is a membrane-associated receptor adaptor with a very short extracellular portion. *KLRC2* encodes NKG2C, an activating cell surface receptor that interacts with HLA-E (the human analogue of mouse Qa-1) on the target cells (*50*). Feature plots showed that *TYROBP* and *KLRC2* expression was also detected in MAIT, Tcm XCL1^+^ and Stem-like cell clusters (Fig. 2F). However, these three clusters could be distinguished from Treg clusters by their expression of *CD27,* with the MAIT cluster additionally characterized by *KLRB1*^hi^*TRAV1-2*^+^*TRAJ33/20/12^+^* phenotype and the Stem-like cell cluster by *CD28* expression, whereas the Treg clusters were *CD27*^−^*CD28*^−^ (Fig. 2B). Thus, among conventional CD8^+^ T cells, Tregs can be defined as *TYROBP^+^KLRC2^+^CD27*^−^ cells. Taken together, our results demonstrate that, in the compartment of conventional CD8^+^ T cells, Tregs can be effectively identified as NKG2C^+^CD27^−^ cells in flow cytometric analysis. A signature consisting of *TYROBP* and *KLRC2* co-expression and the absence of *CD27* serves as a robust classifier for distinguishing CD8^+^ Tregs from other CD8^+^ T cell subpopulations in single-cell transcriptomic data.

### Distinct activation states and gene profiles identify CD8^+^ Treg subsets

Differential gene expression analysis showed that *PTGDS, ZNF683, FGFBP2, CD226 and CD52* were the top upregulated genes in CD8^+^ Treg2 compared to CD8^+^ Treg1, ranked by adjusted *p*-value (Fig. 3A). We generated violin plots for the top 5 upregulated differentially expressed (DE) genes in CD8^+^ Treg2, as well as central memory T cell marker gene *LEF1*. The marked increase in *PTGDS* levels was unique to CD8^+^ Treg2 cells (Fig. 2D and Fig. 3A-C). *PTGDS* encodes prostaglandin D2 (PGD_2_) synthase, a secreted enzyme that catalyzes conversion of prostaglandin H2 (PGH2) to PGD_2_(*51*). PGDis an inflammation mediator (*52*), which regulates immune responses via interaction with its receptors DP1 and DP2 (CRTH2) expressed on immune cells. The transcription factor ZNF683 is also known as Hobit, which is a key regulator of tissue-resident lymphocytes (*53*) and effector T cell differentiation (*54*). *FGFBP2* encodes fibroblast growth factor binding protein 2 (FGF-BP2), which acts as a carrier for FGFs and promotes FGF signaling (*55*). CD226, a cell surface antigen, is a crucial co-stimulatory receptor that amplifies T cell activation (*56*). Soluble CD52 binds inhibitory receptor Siglec-10 on the surface of other immune cells and suppresses T cell proliferation (*57*). Pathway analysis of the DE genes between Treg2 and Treg1 revealed enhanced activity in prostaglandin signaling, fatty acid and lipid metabolism, and T cell and leukocyte activation pathways (Fig. 3D). These findings demonstrate that CD8^+^ Treg2 cells represent a more activated Treg state than CD8^+^ Treg1 cells.

**Fig. 3.**
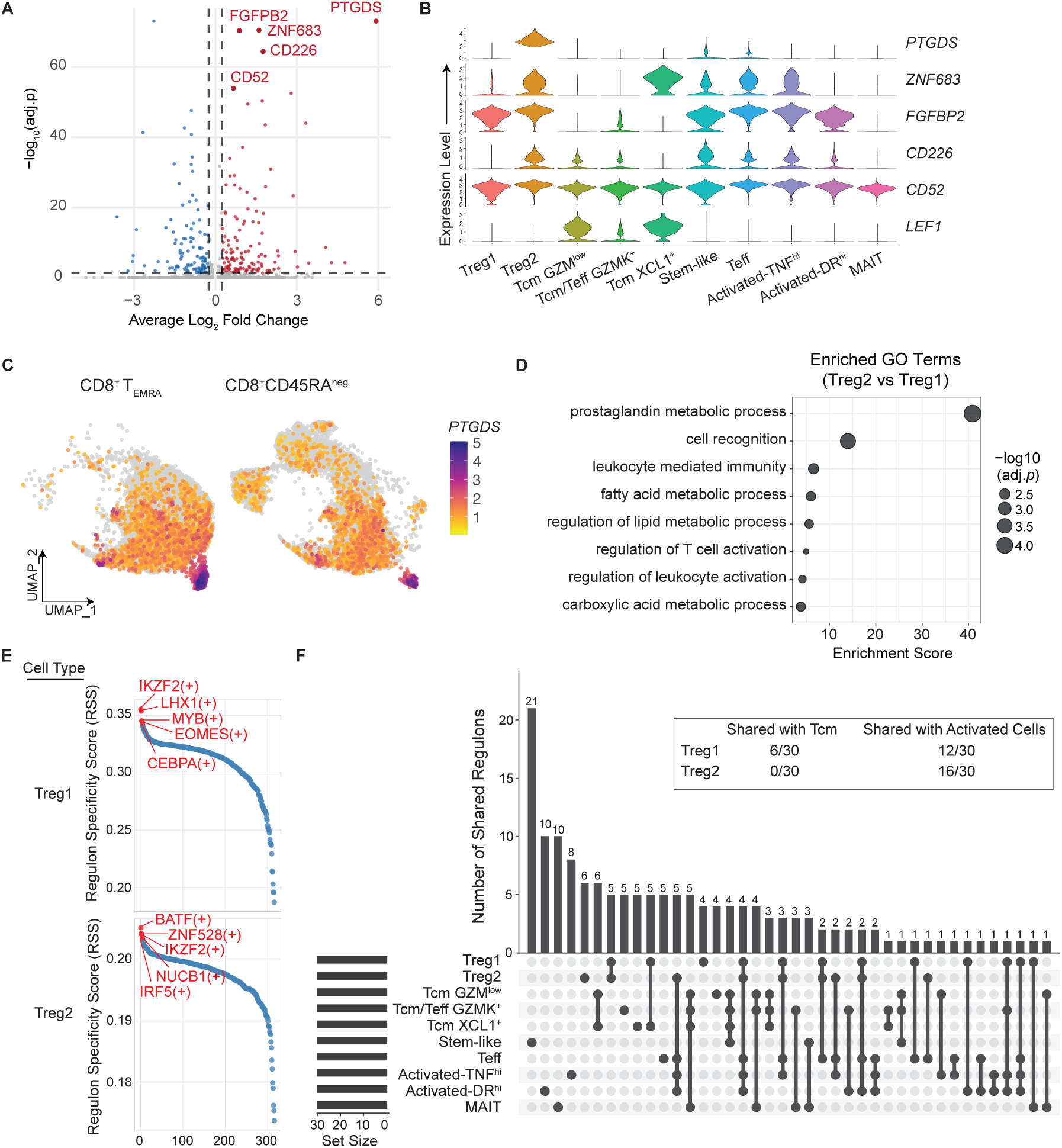
*PTGDS* (encodes prostaglandin D2 synthase) and *CD226*: key markers distinguishing CD8^+^ Treg2 from CD8^+^ Treg1. CITE-seq of CD8^+^ Tand CD45RA^neg^ cells (continued). (**A**) Volcano plot showing the top 5 upregulated genes in CD8^+^ Treg2 versus Treg1 cells. (**B**) Violin plots showing the expression of the top 5 upregulated genes in CD8^+^ Treg2 and Tcm-marker gene *LEF1*. (**C**) UMAP plot of merged CD8^+^ Tand CD45RA^neg^ cells showing the expression of *PTGDS*. (**D**) GO pathway enrichment analysis of the differentially expressed (DE) genes between Treg2 versus Treg1 cells. (**E** & **F**) Gene regulatory network (GRN) analysis of the marker genes of CD8^+^ T cell subsets, performed using pySCENIC. (**E**) The top 5 transcription factor (TF) regulons specific to Treg1 (**upper**) and Treg2 (**lower**) cells. (**F**) UpSet plot showing shared TF regulons across various CD8^+^ T subsets. The **Inset** depicting the number of shared regulons between the top 30 Treg1- and Treg2-specific TF regulons and those of central memory (Tcm) and activated T cells. Tcm cells represent the combined Tcm GZM^low^, Tcm GZMK^+^ and Tcm XCL1^+^ populations. Activated T cells represent the combined Activated-DR^hi^ and Activated-TNF^hi^ populations.

To further explore our finding that CD8^+^ Treg2 cells are more activated than CD8^+^ Treg1 cells, we performed gene regulatory network (GRN) analysis with SCENIC (*58*) to identify most active transcription factor (TF) regulons across the ten CD8^+^ T cell subsets. The top five regulons in CD8^+^ Treg1 were IKZF2, LHX1, MYB, EOMES and CEBPA while those in CD8^+^ Treg2 were BATF, ZNF528, IKZF2, NUCB1 and IRF5 (Fig. 3E). Among them, IKZF2, LHX1, MYB and EOMES are developmental/lineage TFs and BATF, CEBPA and IRF5 are TFs that regulate immune/inflammatory responses (*59–62*). Of the top 30 regulons identified in each of the ten CD8^+^ T cell subsets, CD8^+^ Treg1 shared six regulons with Tcm cells and 12 with activated T cells. In contrast, CD8^+^ Treg2 shared 16 regulons with activated T cells, with no overlap with Tcm cells (Fig. 3F). Thus, our GRN analysis results also indicate that CD8^+^ Treg2 represents a more activated Treg state than CD8^+^ Treg1.

In summary, CD8^+^ Treg2 cells exhibit higher expression of genes related to immune activation and inflammation, such as *PTGDS*, *ZNF683*, *FGFBP2*, *CD226*, and *CD52*, compared to CD8^+^ Treg1 cells. Pathway and gene regulatory network analyses further reveal that the CD8^+^ Treg2 is a more activated Treg subset, with enhanced signaling and greater overlap with activated T cells, whereas CD8^+^ Treg1 shares features with central memory T cells.

### CD8^+^ Tregs exhibit high clonality and share TCRs with diverse CD8^+^ T cell subpopulations

We conducted paired scTCR-seq on sorted CD8^+^ Tand CD45RA^neg^ cells to investigate the characteristics and diversity of the TCR repertoires of CD8^+^ Tregs. Among the 22,651 high-quality single cells (11,181 of Tcells and 11,490 of CD45RA^neg^ cells), 18,633 cells (82.3%) yielded productive paired TCR αβ sequences. Among these, 4,196 were Tregs, comprising 3,700 Treg1 and 496 Treg2 cells. None of these 4,196 cells expressed the invariant NKT (iNKT)-specific TCR Vα24-Jα18 (*63*). Both CD8^+^ Treg1 and Treg2 cells exhibited high clonality (Fig. 4A and fig. S4A). Quantification of TCR repertoire diversity using the Shannon index, which accounts for both clonal richness and evenness, revealed comparably low diversity in both CD8^+^ Treg1 and Treg2 cells, indicating similarly restricted repertoires across these two Treg subsets (Fig. 4B). Inter-subset sharing of TCRαβ clonotypes was interrogated using an UpSet plot, where a clonotype was defined as a unique combination of TCRα and TCR β chain CDR3 sequences. Both CD8^+^ Treg1 and Treg2 cells shared TCRs with other CD8^+^ T cell subsets (Fig. 4C). To visualize the distribution of a shared TCRαβ clonotype among CD8^+^ T cell subsets, we color coded (blue dots) T cells expressing that clonotype on the UMAP plot of merged CD8^+^ Tand CD45RA^neg^ cells from each donor. Representative plots revealed that a highly expanded clonotype could be detected in either the Treg1 cluster, the Treg2 cluster or both, while also being present in other CD8^+^ T cell subtypes, such as the Stem-like and Teff clusters (Fig. 4D). Similar TCR-sharing patterns were observed across different individuals (fig. S4B). Morisita index analysis further quantified these observations, showing moderate overlap between Treg1 and Treg2 (Cδ = 0.470). In our CITE-seq analysis, the Tcm XCL1^+^ cluster expressed Treg marker genes (*TYROBP^+^KLRC2^+^IKZF2^+^KIR^+^*) but, unlike CD27^−^ Treg clusters, was CD27^+^ (Fig. 2B and 2F); in the Morisita index analysis, the cluster exhibited no overlap with either Treg1 (Cδ = 0.000) or Treg2 (Cδ = 0.000) (Fig. 4E). Thus, the absence of CD27 expression is critical for differentiating Treg1 and Treg2 from Tcm XCL1^+^ cells. Interestingly, while Treg2 exhibited moderate overlap with Stem-like (Cδ = 0.532), Teff (Cδ = 0.563) and Activated-TNF^hi^ (Cδ = 0.581) cells, Treg1 showed very low overlap with Stem-like (Cδ = 0.192), Teff (Cδ = 0.060) and Activated-TNF^hi^ (Cδ = 0.212) cells (Fig. 4E). These findings suggest that Treg1 and Treg2 may follow distinct developmental routes during the development of CD8^+^ Tregs.

**Fig. 4.**
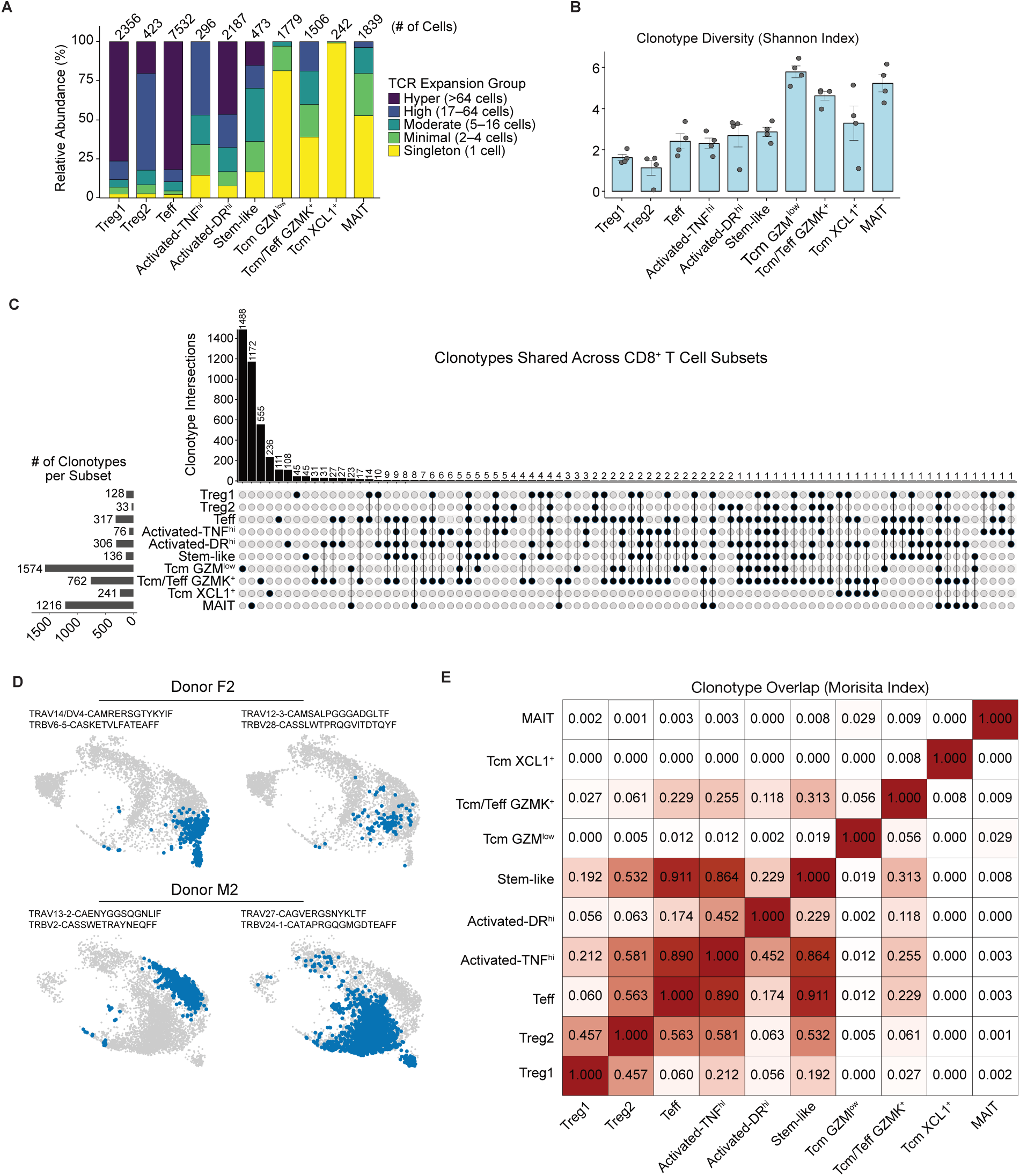
CD8^+^ Tregs exhibit high clonality and share T-cell receptors with diverse CD8^+^ T cell subpopulations. scTCR-seq of CD8^+^ Tand CD45RA^neg^ cells (continued). (**A**) Relative abundance of cells with varying degrees of clonality (singleton, minimal, moderate, high and hyper) within each CD8^+^ T cell subpopulation. The number above each column indicates the number of cells in the corresponding subpopulation. (**B**) Shannon diversity index for TCRαβ clonotypes across CD8^+^ T cell subpopulations (*n* = 4, mean ± s.d.). (**C**) UpSet plot showing shared TCR αβ clonotypes across CD8^+^ T cell subpopulations. (**D**) UMAP plots of merged CD8^+^ Tand CD45RA^neg^ cells of donor F2 (upper panel) and donor M2 (lower panel). Four representative TCRαβ clonotypes are depicted, each of which shared between Treg1 and/or Treg2 cells and other CD8^+^ T cell subpopulations. Blue dots represent individual cells expressing the TCRαβ clonotype indicated above the corresponding UMAP plot. (**E**) Morisita index analysis of TCRαβ clonotype overlap across CD8^+^ T cell subpopulations.

### NKG2C^+^CD27^−^ defines a CD8^+^ Treg-specific phenotype, while CD226 differentiates Treg1 and Treg2 cells

To validate the novel surface markers of CD8^+^ Tregs we identified through our CITE-seq analysis, we developed a 24-antibody panel for flow cytometric analysis (table S3), enabling high-resolution characterization of CD8^+^ T cells, with a focus on CD8^+^ Tregs. We analyzed PBMCs from eight healthy donors. Following the exclusion of γδT cells using anti-γδTCR and MAIT cells using anti-TCR Vα7.2 within the CD3^+^ T cell compartment, we gated on CD8^+^CD4^−^ cells to isolate the conventional CD8^+^ T (CD8^+^ Tcon) cell compartment. Within this population, we observed that the surface phenotype of NKG2C^+^CD27^−^ defines a distinct cell population—CD8^+^ Tregs (Fig. 5A). Specifically, among CD3^+^ Tcon cells, the NKG2C^+^CD27^−^ population was found exclusively within the CD8^+^ T cell compartment, with only minimal numbers of NKG2C^+^CD27^−^ cells detected in the CD4^+^ T cell compartment (Fig. 5A-B). CD8^+^ Tregs constitute 2.7% (± 3.3%, s.d.) of total CD8^+^ Tcon cells (Fig. 5B). Of note, 11.0% (± 16.1%, s.d.) of total γδT cells and 1.4% (± 1.9%, s.d.) of CD8^+^ MAIT cells displayed the phenotype of NKG2C^+^CD27^−^ (fig. S5A-D). Therefore, exclusion of γ δ T and CD8^+^ MAIT cells is essential for accurate analysis of NKG2C^+^CD27 CD8^+^ Tregs.

**Fig. 5.**
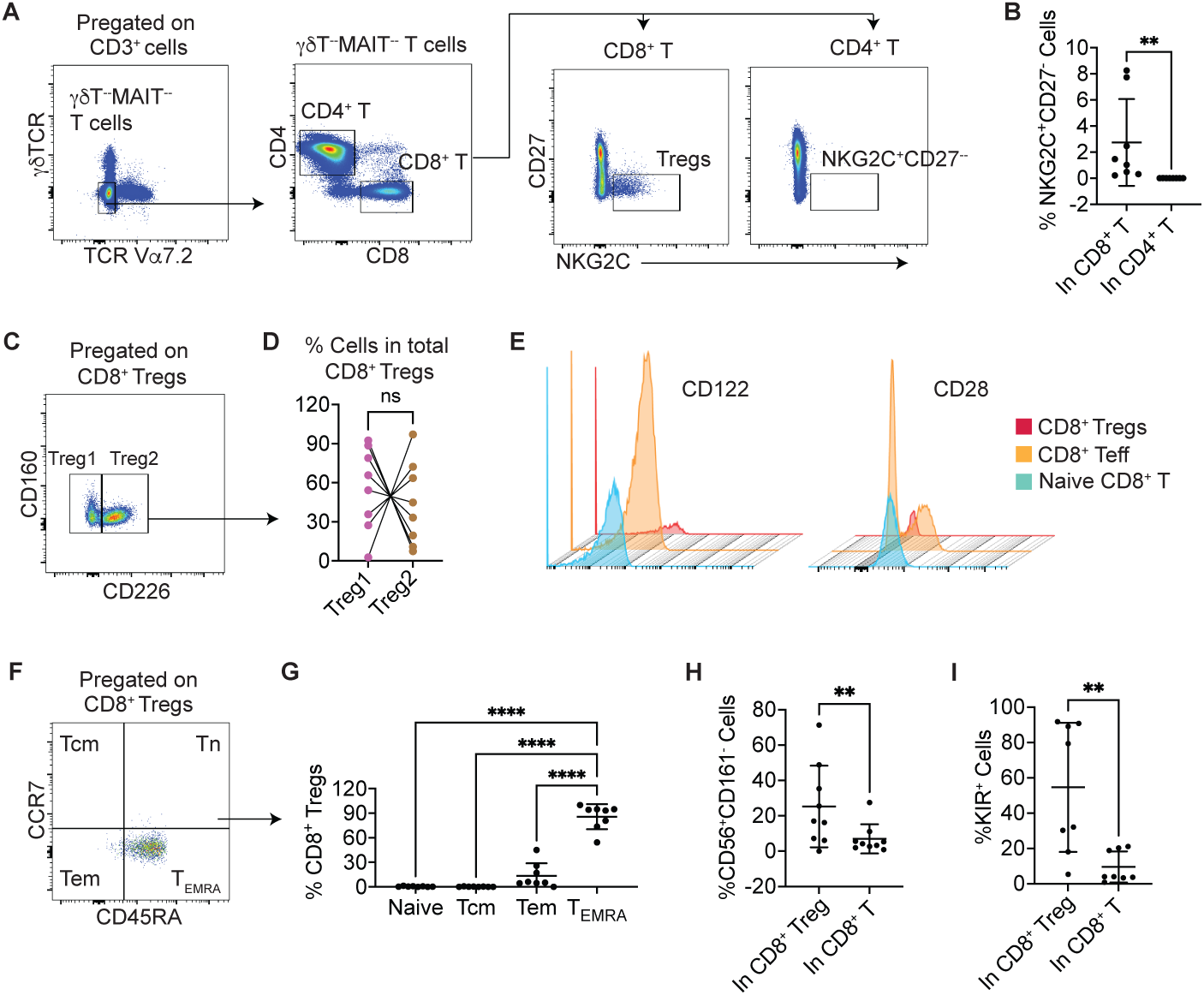
Flow cytometric validation: NKG2C^+^CD27^−^ defines a CD8^+^ Treg-specific phenotype, while CD226 differentiates Treg1 and Treg2. PBMCs from HCs were analyzed using a CyTek® Aurora flow cytometer with a 24-antibody panel. (**A**) Representative plots illustrating the gating strategy for NKG2C^+^CD27^−^ CD8^+^ Tregs and showing that NKG2C^+^CD27^−^ cells were largely absent from the CD4^+^ T cell compartment. (**B**) CD8^+^ Treg frequencies in CD8^+^ T cells and NKG2C^+^CD27^−^ cell frequencies in CD4^+^ T cells. Wilcoxon matched-pairs signed-rank test, \*\**p* < 0.001 (*n* = 8, mean ± s.d.). (**C**) Representative plot illustrating that CD226 expression levels distinguish Treg1 (low) and Treg2 (high). (**D**) Frequencies of Treg1 and Treg2 within the CD8^+^ Treg compartment. Each line connects the two samples from the same individual. Wilcoxon matched-pairs signed-rank test; ns, not significant (*n* = 8). (**E**) Representative histogram plots showing CD8^+^ Tregs were CD122^hi^CD28^−^. (**F**) Representative plot illustrating CD45RA and CCR7 expression in CD8^+^ Tregs, markers that delineate T cells into four subsets: Tn, Tcm, Tem and Tcells. (**G**) Frequencies of Tn, Tcm, Tem and Tcells among CD8^+^ Tregs. Ordinary one-way ANOVA, Dunnett’s multiple comparisons test, **** *p* < 0.0001 (*n* = 8, mean ± s.d.). (**H**) Frequencies of CD56^+^CD161^−^ cells in CD8^+^ Tregs and total conventional CD8^+^ T cells, respectively. Wilcoxon matched-pairs signed-rank test, \*\**p* =<0.001 (*n* = 8, mean ± s.d.). (**I**) Frequencies of KIR^+^ cells in CD8^+^ Tregs and total conventional CD8^+^ T cells, respectively. Wilcoxon matched-pairs signed-rank test, \*\**p* =<0.001 (*n* = 8, mean ± s.d.).

Further subset analysis revealed that CD226 expression divided the CD8^+^ Treg population into two subsets—Treg1 (CD226^−^) and Treg2 (CD226^+^) (Fig. 5C). Within the Treg population, there was no difference in the proportions of these two subsets (Fig. 5D). Consistent with our CITE-seq results, CD8^+^ Tregs exhibited a CD122^+^CD28^−^CCR7^−^ surface phenotype (Fig. 5E-F) and 85.8% (±15.4%, s.d.) of CD8^+^ Tregs were Tcells (Fig. 5G), further supporting the validity of the markers we identified through our high-dimensional profiling approach. Both CD8^+^CD56^+^ and CD8^+^KIR^+^ T cells were enriched in the NKG2C^+^CD27^−^ CD8^+^ Treg compartment (Fig. 5H-I).

In summary, we validated the novel surface markers for CD8^+^ Tregs identified through our CITE-seq analysis using flow cytometry. Based on this validation, we defined CD8^+^ Tregs as CD3^+^γδTCR^−^Vα7.2^−^CD8^+^NKG2C^+^CD27^−^ cells. Furthermore, we found that CD226 expression partitioned CD8^+^ Tregs into two distinct subsets: Treg1 (CD226^−^) and Treg 2 (CD226^+^). Notably, the large majority of CD8^+^ Tregs fell within the Tcell compartment.

### Functional analysis of CD8^+^ Treg-mediated suppression in the mixed lymphocyte reaction

To assess the suppressive function of CD8^+^ Tregs, we performed a mixed lymphocyte reaction (MLR) assay (Fig. 6A). Mitomycin C (MMC)-treated autologous PBMCs were mixed with equal number of allogeneic PBMCs and used as stimulators/antigen presenting cells (APCs), and CellTrace Violet (CTV)-labeled CD8^+^ Treg-depleted CD3^+^ T cells served as responders/target cells (CTV-labeled target cells). Total CD8^+^ Tregs (tTreg) (CD3^+^γδTCR^−^Vα7.2^−^CD8^+^NKG2C^+^CD27^−^), Treg1 (CD226^−^ Tregs), Treg2 (CD226^+^ Tregs) and control CD8^+^ T cells (CD3^+^γδTCR^−^Vα7.2^−^CD8^+^NKG2C^−^) as effector cells were isolated by cell sorting (table S4 and fig. S5E) and co-cultured with target cells. Effector and target cells were co-cultured at a 1:4 ratio (1 Treg to 4 target cells) with CD8^+^ Tregs or control effector cells comprising 20% of the total T cell population. After four days, T cells were analyzed by flow cytometry using an antibody panel recognizing CD3, CD4, CD8, NKG2C, CD27, CD226, CD56, CD161, KIR2DL2/L3/S2, and KIR3DL1. Four days after co-culture of targets with APCs, distinct CTV^hi^ peak and CTV^-/low^ populations were observed in both CD4^+^ and CD8^+^ T cell populations, indicating that proliferation was induced in the MLR assay (fig. S5F). CTV dilution analysis on gated CD3^+^CD4^+^ target cells showed no differences among co-culture conditions containing control CD8^+^ T cells, tTregs, Treg1 or Treg2 (fig. S5G). We then examined the frequency of T cell subpopulations and found that the frequencies of Tregs in co-culture remained stable across all conditions (Fig. 6B, Left), indicating that any observed suppressive effects were not attributed to differential expansion or loss of the regulatory populations themselves. The frequencies of CD8⁺ T cells were significantly reduced in co-cultures containing tTreg, Treg1, or Treg2 cells, relative to control co-cultures containing non-regulatory control CD8⁺ T cells (Fig. 6B, Middle). This reduction suggests active suppression of CD8⁺ responder T cells by all three Treg populations. In contrast, no differences in CD4⁺ T cell frequencies were detected across any of the co-culture conditions (Fig. 6B, Right). Human CD8^+^ Tregs have been reported to suppress both CD4^+^ and CD8^+^ T cells (*18–20, 26, 64*). Therefore, it was unexpected that, in our MLR assay, CD8^+^ Tregs selectively reduced CD8⁺ T cell frequencies without affecting CD4⁺ T cell frequencies. These observations suggest that the suppressive activity of these Treg subsets may be selective for the CD8⁺ compartment under the assay conditions tested.

**Fig. 6.**
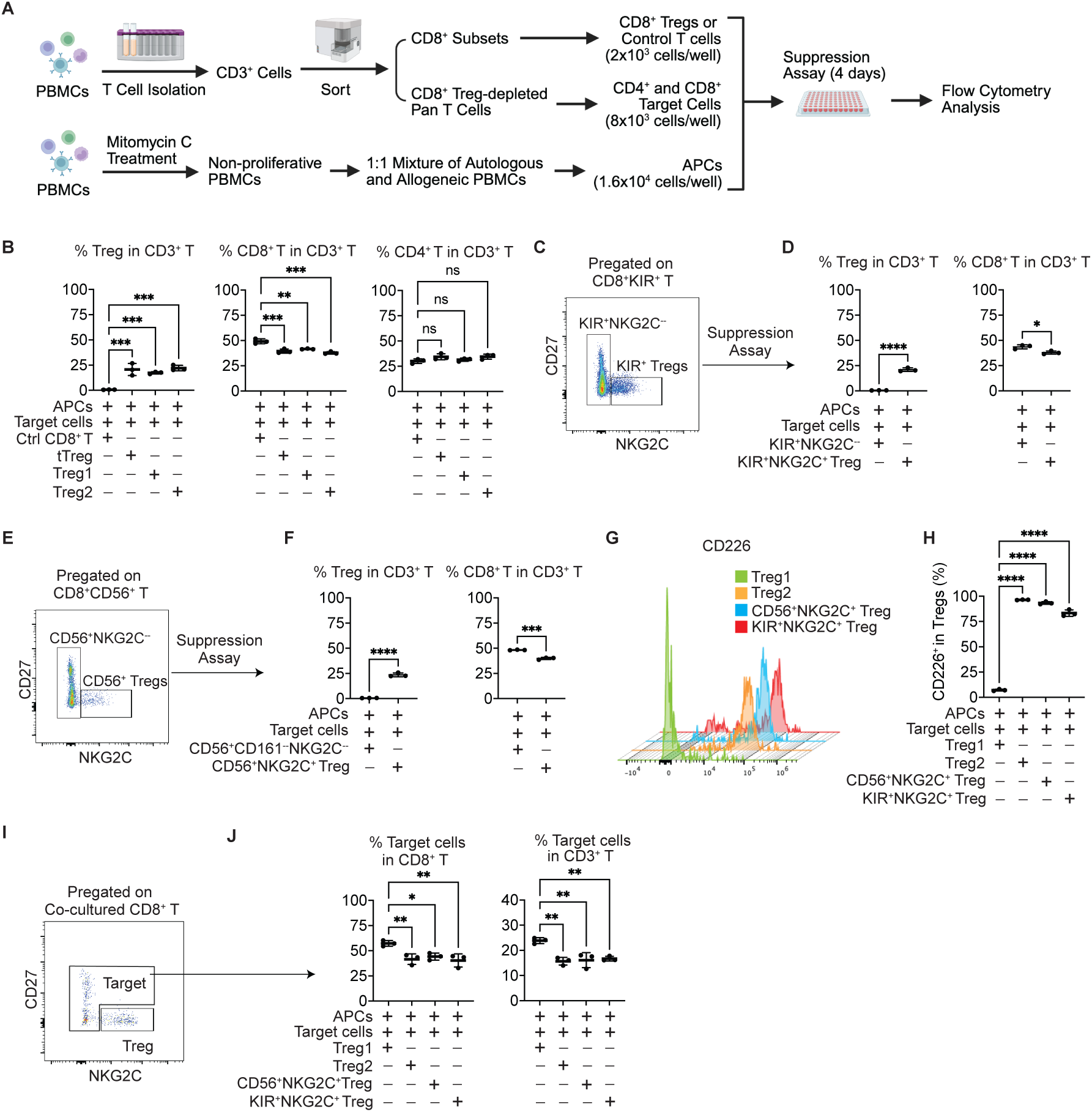
CD8^+^ Tregs suppress CD8^+^ T cells *in vitro*, with Treg2 showing stronger suppressive activity than Treg1. (**A**) Scheme of CD8^+^ Treg-mediated suppression assay (created using BioRender.com). NKG2C^+^CD27^−^ total CD8^+^ Tregs (tTreg), Treg1, Treg2, CD8^+^KIR^+^NKG2C^+^CD27^−^ (KIR^+^NKG2C^+^ Treg) and CD8^+^CD56^+^CD161^−^NKG2C^+^CD27^−^ Treg (CD56^+^NKG2C^+^ Treg) were sorted as effector (Treg) cells, while NKG2C^−^, KIR^+^NKG2C^−^ and CD56^+^CD161^−^NKG2C^−^ CD8^+^ T cells were sorted as control effector cells. tTreg-depleted T cells were sorted as target cells. In **B, D, F, H,** and **J**, co-cultured cells were harvested on day 4 and analyzed by flow cytometry, and the x-axis indicates the co-culture conditions. (**B**) Left, frequencies of NKG2C^+^CD27^−^ CD8^+^ Tregs among total T cells; middle, frequencies of CD8^+^ T cells among total T cells; right, frequencies of CD4^+^ T cells among total T cells. Ordinary one-way ANOVA, Dunnett’s multiple comparisons test, ** *p* < 0.01, *** *p* < 0.001 (*n* = 3, mean ± s.d.). (**C**) Differential expression of NKG2C and CD27 on CD8^+^KIR^+^ T cells divided the population into KIR^+^NKG2C^+^ Treg and KIR^+^NKG2C^−^ subpopulations. (**D**) Left panel, frequencies of NKG2C^+^CD27^−^ CD8^+^ Tregs among total T cells; left panel, frequencies of CD8^+^ T cells among total T cells; right panel. Unpaired *t*-test, * *p* < 0.05, **** *p* < 0.0001. (**E**) Differential expression of NKG2C and CD27 on CD8^+^CD56^+^CD161^−^ (CD8^+^CD56^+^) T cells divided the population into CD56^+^NKG2C^+^ Treg and CD56^+^CD161^−^NKG2C^−^ subpopulations. (**F**) Left panel, frequencies of NKG2C^+^CD27^−^ CD8^+^ Tregs among total T cells; left panel, frequencies of CD8^+^ T cells among total T cells; right panel. Unpaired *t*-test, *** *p* < 0.001, **** *p* < 0.0001. (**G**) On day 4, histogram depicting the expression of CD226 on pregated Treg1, Treg2, CD56^+^NKG2C^+^ Tregs and KIR^+^NKG2C^+^ Tregs of the co-culture. (**H**) Frequencies of CD226^+^ cells among Treg1, Treg2, CD56^+^NKG2C^+^ Tregs and KIR^+^NKG2C^+^ Tregs. (**I**) Representative plot showing that differential expression of NKG2C and CD27 distinguished Tregs from target cells in a Treg and target co-culture system. (**J**) Left, frequencies of target cells among CD8^+^ T cells; right, frequencies of target cells among total T cells. Ordinary one-way ANOVA, Dunnett’s multiple comparisons test, * *p* < 0.05, ** *p* < 0.01.

To compare the suppressive activity of CD8^+^KIR^+^NKG2C^+^CD27^−^ (KIR^+^NKG2C^+^ Treg) versus CD8^+^KIR^+^NKG2C^−^ (KIR^+^NKG2C^−^) cells, these two subsets were isolated by cell sorting (Fig. 6C) and then evaluated in the MLR assay. After four days, T cells were analyzed by flow cytometry. CTV dilution analysis on gated CD3^+^CD4^+^ responder cells revealed no differences between the two co-culture conditions (fig. S5H). The frequencies of KIR^+^NKG2C^+^ Tregs in co-culture remained comparable to tTreg, Treg1 and Treg2 (Fig. 6D, Left), while the frequency of CD8⁺ T cells was reduced in the co-culture containing KIR^+^NKG2C^+^ Tregs relative to the control co-culture containing KIR^+^NKG2C^−^ cells (Fig. 6D, Right). Similarly, CD8^+^CD56^+^CD161^−^NKG2C^+^CD27^−^ (CD56^+^NKG2C^+^ Tregs) and CD8^+^CD56^+^CD161^−^NKG2C^−^(CD56^+^CD161^−^NKG2C^−^) cells were isolated and analyzed in the MLR assay (Fig. 6E). CTV dilution analysis on gated CD3^+^CD4^+^ responder cells revealed no differences between the two co-culture conditions (fig. S5I). We found that the frequencies of KIR^+^NKG2C^+^ Tregs in co-culture remained comparable (Fig. 6F, Left), whereas the frequency of CD8⁺ T cells was reduced in the co-culture containing CD56^+^NKG2C^+^ Tregs, relative to the control co-culture containing CD56^+^CD161^−^NKG2C^−^ cells (Fig. 6D, Right). These results demonstrate that the NKG2C^+^CD27^−^ phenotype defines bona fide CD8^+^ Tregs.

Flow cytometric analysis performed after four days of co-culture revealed that the frequencies of CD226^+^ cells in the NKG2C^+^CD27^−^ CD8⁺ Treg compartment were as follows: 7.1% in cultures containing Treg1, 96.4% in cultures containing Treg2, 93.3% in cultures containing CD56^+^NKG2C^+^ Tregs, and 83.3% in cultures containing KIR^+^ Tregs, respectively (Fig. 6G-H). These results demonstrated that Treg1 and Treg2 populations maintained their respective CD226 expression patterns. Moreover, a large majority of CD56^+^ Tregs and KIR^+^ Tregs were Treg2 cells since they expressed CD226. Given that CD8^+^ responders (NKG2C^−^ and NKG2C^+^CD27^+^ cells) could be distinguished from Tregs (NKG2C^+^CD27^−^) based on differential expression of NKG2C and CD27 in co-cultures containing Treg1, Treg2, CD56^+^NKG2C^+^ Treg or KIR^+^NKG2C^+^ Treg cells (Fig. 6I), we assessed the frequencies of CD8^+^ responders under these conditions. We found that the frequencies of CD8^+^ responders—whether measured in total CD8 T cells or in CD3^+^ T cells—were higher in cultures containing Treg1 compared to other conditions that contained Treg2 or were enriched for Treg2 (Fig. 6J). This indicates that Treg2 cells were more effective at suppressing CD8⁺ responder proliferation relative to Treg1, consistent with our CITE-seq findings showing that Treg2 cells represent a more activated regulatory state than Treg1 cells.

In summary, we investigated the suppressive function of various CD8^+^ Treg subsets using tailored MLR assay. We found that total Tregs (tTreg) and Treg1 and Treg2 subsets, as well as CD56^+^NKG2C^+^ and KIR^+^NKG2C^+^ Tregs, all significantly reduced the frequency of CD8^+^ responder T cells, indicating active suppression, whereas CD4^+^ T cell frequencies were unaffected in the MLR assay. The NKG2C^+^CD27^−^ phenotype was identified as a marker for bona fide CD8^+^ Tregs. Treg2 cells (CD226^+^) were found to be more effective at suppressing CD8^+^ responder proliferation than Treg1 cells (CD226^−^). These findings demonstrate that specific CD8^+^ Treg subsets selectively suppress CD8^+^ T cell responses under the assay conditions tested, whereas Treg2 cells represented a more activated and potent regulatory state.

### CD8^+^ Tregs increase with age

To investigate age-associated changes in CD8^+^ Treg populations, we analyzed publicly available CITE-seq and scTCR-seq dataset syn49637038 from PBMCs of 166 healthy donors spanning an age range of 25 to 85 years (*65*). We used antibody-derived tags (ADTs) to enrich for high-quality CD3^+^CD4^−^CD8^+^ (CD8^+^ T) cells (271,875 cells) followed by further enrichment for CD45RA^+^CD27^−^ (CD8^+^ T) cells (45,390 cells) (Fig. 7A). We used CD8^+^ Treg marker genes, including *TYROBP*, *KLRC2*, *IKZF2*, *ICAM1*, *KIR3DL1*, *KIR2DL3*, *KIR2DL1* and *PTGDS* to identify CD8^+^ Treg1 (Cluster 0: 12,753 cells) and Treg2 (Cluster 11: 1,206 cells) clusters in CD8^+^ Tcells (Fig. 7B and fig. S6A-B). To evaluate the reproducibility and biological consistency of the Treg2 vs Treg1 signature between our CITE-seq dataset and this public dataset, we performed a concordance analysis of differentially expressed (DE) genes. DEGs were defined as |log₂ fold change| > 0.50 and FDR < 0.05 in each dataset separately. Of the 95 DEGs identified in the two datasets, 73 (76.8%) showed concordant directional change (*r* = 0.206, *p* < 0.001), which was visualized with a four-quadrant scatter plot (Fig. 7C). Top concordant upregulated genes in Treg2 included *PTGDS*. This concordance confirms the robustness of the Treg1 and Treg2 transcriptional signatures across datasets. The frequencies of these Treg1 and Treg2 cells did not vary by sex (fig. S6C), whereas the frequencies of CD8^+^ TTreg1 and Treg2 cells increased with age (Fig. 7D). Furthermore, within the CD8^+^ Tcompartment, Treg2 but not Treg1 frequencies increased with age (Fig. 7E).

**Fig. 7.**
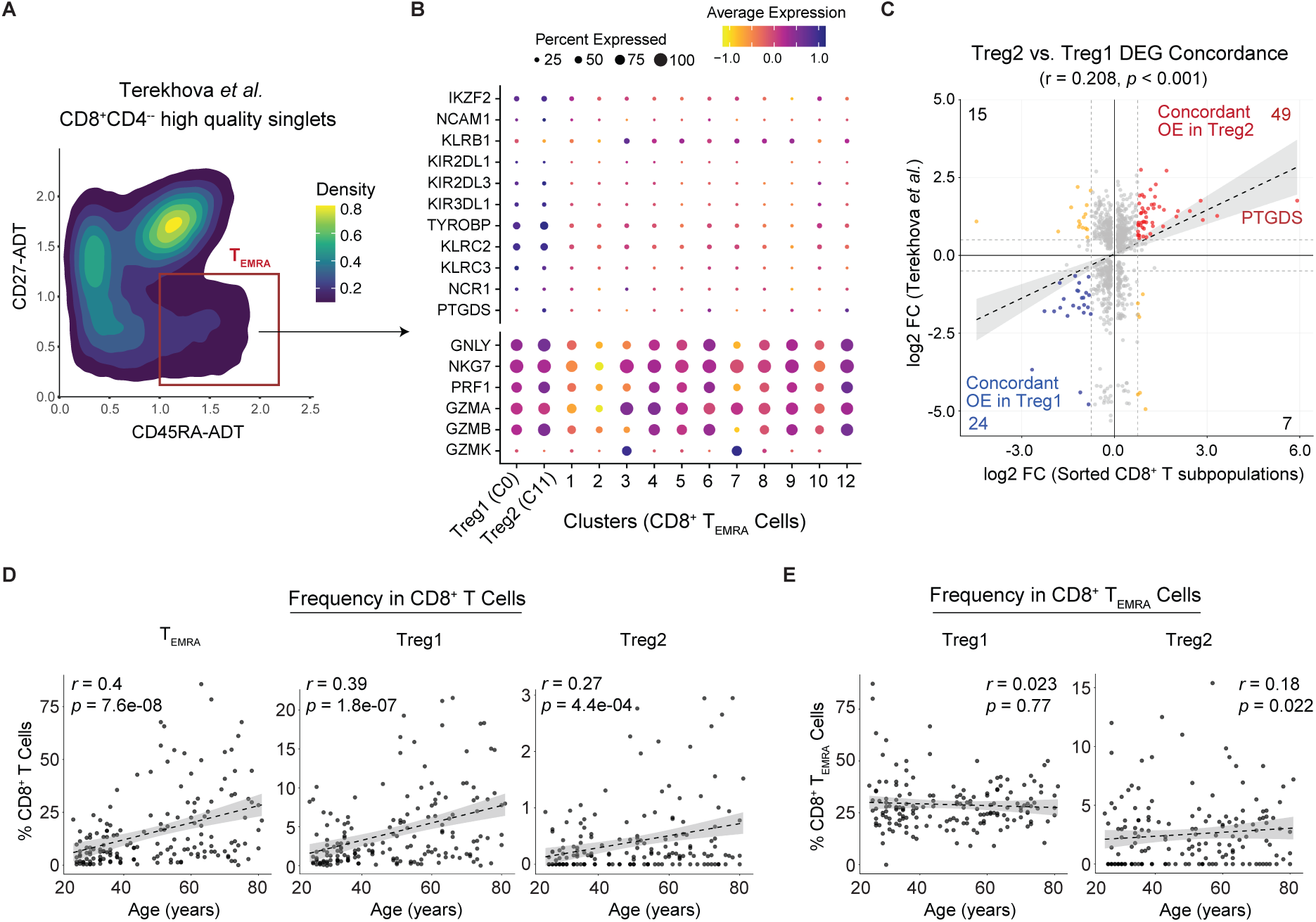
The frequency of CD8^+^ Treg2 increases with aging among CD8^+^ Tcells. CITE-seq and scTCR-seq expression profiles of 166 healthy donors aged 25-85 years old were extracted from the public scRNA-seq dataset syn49637038. (**A**) ADT biaxial plot: after excluding MAIT, γδ T and CD4^+^ T cells, high-quality CD8^+^ T cell singlets were used to generate an ADT biaxial plot showing CD45RA versus CD27 expression. From this plot, CD45RA^+^CD27^−^ CD8^+^ Tsinglets were extracted for the downstream analysis. (**B**) Dot plot showing the expression of CD8^+^ Treg-associated and cytotoxicity-related genes across CD8^+^ Tcell clusters, and identification of Treg1 and Treg2 clusters with *TYROBP*, *KLRC2* and *PTGDS* expression. C0 and C11 indicate cluster numbers. (**C**) Concordance analysis was performed between DE genes (Log2 FC > 0.50 or < −0.50, adj.*p* < 0.05) identified from two comparisons: (1) Treg2 versus Treg 1 of the sorted CD8^+^ Tand CD45RA^neg^ cell dataset, and (2) Treg2 versus Treg1 identified within the CD45RA^+^CD27^−^ CD8^+^ Tsinglets of the public dataset. Among 869 genes analyzed, 95 DE genes were identified (Log2 FC > 0.50 or < −0.50, adj.*p* < 0.05). Red dots, genes concordantly overexpressed in Treg 2 (*n* = 49); blue dots, genes concordantly overexpressed in Treg 1 (*n* = 24); yellow dots, discordantly expressed genes (*n* = 22). OE, overexpression. (**D**) The frequency of CD8^+^ T, Treg1 and Treg2 increased with aging among total CD8^+^ T cells. (**E**) The frequency of Treg2 increased with aging among CD8^+^ Tcells, while the frequency of Treg1 remained stable.

## DISCUSSION

In this study, we identified two transcriptionally distinct CD8^+^ Treg subsets, designated Treg1 and Treg2. While both subsets share a core phenotype of CD3^+^γδTCR^−^Vα7.2^−^CD8^+^NKG2C^+^CD27^−^, they are distinguished by differential expression of surface antigen CD226, as well as by differential expression of intracellular enzyme prostaglandin D2 synthase (PTGDS). Specifically, Treg1 is CD226^−^PTGDS^−/low^, whereas Treg2 is CD226^+^PTGDS^hi^. Differential gene expression (DEG), gene regulatory network (GRN), and mixed lymphocyte reaction (MLR) analyses collectively revealed that Treg2 represents a more activated Treg state than Treg1. CD8^+^ Tregs are highly clonal, and Treg1 and Treg2 may follow distinct developmental routes. Almost all CD8^+^ Tregs reside within the Tcompartment. Within the total CD8^+^ T cell population, the frequencies of both CD45RA^+^CD27^−^ Treg1 and Treg2 cells increase with age; however, within the Tcompartment, only Treg2 frequency increases with age, whereas Treg1 remains stable.

The field of CD8^+^ Treg research has been complicated by the description of various CD8^+^ Treg subsets defined by disparate and often non-overlapping cell surface markers, including CD8^+^Ly49^+^, CD8^+^CD56^+^CD161^−^, CD8^+^KIR^+^, CD8^+^CD122^+^, CD8^+^CD28^−/low^ and others across mouse and human studies (*10, 13, 14, 18–22*). The lack of consensus on a unified phenotypic signature has led to several fundamental challenges in understanding this important population. First, direct comparison of findings across studies is difficult, as different groups have isolated CD8^+^ Treg using different marker combinations, potentially capturing distinct or only partially overlapping populations. Second, some of these markers, such as Ly49 receptors and KIRs, belong to families of proteins with stochastic expression (*15, 17*), meaning that the identity of CD8^+^Ly49^+^ and CD8^+^KIR^+^ Tregs may shift under different physiological or genetic conditions, and the common practice of using multiple Ly49 or KIR proteins together for identification further complicates reproducible isolation and functional characterization. Third, the absence of a reliable core signature raises the possibility that many so-called CD8^+^ Treg-enriched populations are heterogeneous mixtures containing both genuine regulatory cells and conventional CD8^+^ T cells lacking regulatory capacity. Without a consensus phenotype, cross-study comparison remains problematic, and progress toward defining CD8^+^ Treg ontogeny and transcriptional regulation is substantially hindered. Consequently, translating CD8^+^ Treg biology into therapeutic applications has remained elusive.

In this study, we addressed these long-standing challenges by identifying a core phenotypic signature—CD3^+^γδTCR^−^Vα7.2^−^CD8^+^NKG2C^+^CD27^−^—that consolidates these previously described CD8^+^ Treg subsets into a unified cellular identity. The advantage of this approach is twofold. First, by defining the common denominator among diverse reported CD8^+^ Treg populations, our core markers enable the selective enrichment of genuine regulatory CD8^+^ T cells while excluding the substantial contamination of non-Treg cells that hampered previous isolation strategies. Indeed, our CITE-seq and functional analyses reveal that while those subsets contained a fraction of true CD8^+^ Tregs, the majority of cells within those populations were non-regulatory conventional CD8^+^ T cells. Second, the addition of secondary discriminatory markers—specifically CD226 and PTGDS—further refines this core identity to distinguish functionally distinct Treg subpopulations (Treg1 vs. Treg2), allowing for more precise functional stratification. Thus, our core marker panel provides a much-needed standardized framework for CD8^+^ Treg identification, enabling reproducible isolation, cross-study comparability, and a clearer roadmap for harnessing CD8^+^ Tregs for therapeutic purposes.

A major conceptual advance emerging from this study is the recharacterization of CD8^+^ Tcells. In chronic infections such as human immunodeficiency virus (HIV) infection and in autoimmune disorders, excessive accumulation of CD8^+^ Tcells can lead to tissue damage due to their pro-inflammatory cytokine production and cytotoxic activity (*34–36*). In recent years, accumulating studies indicate that CD8^+^ Tcells are also implicated in neurodegeneration and neuroinflammation such as Alzheimer’s disease (AD) (*37–43*). Given the historical view of CD8^+^ Tcells, which represent cytotoxicity and inflammation, it is believed that AD-associated CD8^+^ Tcells fuel neuroinflammation and neurodegeneration in AD patients (*38*). However, preclinical mouse models of AD suggest a more complex role for CD8^+^ T cells in disease pathology. These studies have shown that brain-infiltrating CD8^+^ T cells can either function as CTLs, exacerbating neurodegeneration (*66, 67*), or serve as regulatory T cells (Tregs), thereby helping in reducing amyloid plaque deposition and cognitive decline (*68*). Our results demonstrate that up to 35% of CD8^+^ Tcells exhibit a regulatory transcriptional and functional profile, whereas the vast majority of CD8^+^ Tregs—approximately 85%—are Tcells. Thus, in line with observations in preclinical mouse models, AD-associated CD8^+^ Tcells may include both CTLs and Tregs. Further investigation is needed to delineate the subtypes of AD-associated CD8^+^ Tcells to define their roles in AD pathogenesis.

The finding that a significant portion of CD8^+^ Tcells are Tregs substantially changes our understanding of Tcell biology: these cells are not a uniformly pathogenic population but rather a functionally diverse compartment encompassing both proinflammatory and immunosuppressive subsets, with the latter representing a considerable proportion. The clinical implications of identifying CD8^+^ Tregs within the compartment of CD8^+^ Tcells is twofold. First, disease-associated expansion of CD8^+^ Tcells cannot be interpreted solely as a marker of active inflammation or malignancy, as a substantial regulatory component may be simultaneously mobilized. Second, therapeutic strategies aimed at depleting CD8^+^ Tcells in autoimmune or inflammatory diseases risk eliminating a large reservoir of regulatory cells, potentially exacerbating rather than ameliorating disease. Instead, selectively modulating the balance between proinflammatory CD8^+^ Tcells and CD8^+^ Tregs offers a more targeted therapeutic approach.

A common feature of CD8^+^ Tregs, iNKT cells and NK-like CD8⁺ T cells is that all three express multiple NK cell markers. However, our scTCR-seq data demonstrated that NKG2C^+^CD27^−^ CD8^+^ Tregs are not iNKT cells since they did not express iNKT TCR Vα24-Jα18. CD8^+^ Tregs are also distinct from the broad category of NK-like CD8⁺ T cells—comprising conventional CD8⁺ T cells that upregulate NK receptors such as CD56, KIRs and NKG2A without a defined regulatory function (*69*)—whereas CD8^+^ Tregs possess clear immunosuppressive capacity. By the broader definition of NK-like CD8⁺ T cells, NKG2C^+^CD27^−^ CD8⁺ Tregs, CD8^+^CD56^+^ cells and CD8^+^KIR^+^ cells are all considered NK-like CD8⁺ T cells. However, our data demonstrate that within the CD8^+^CD56^+^ and CD8^+^KIR^+^ populations, only cells displaying the NKG2C^+^CD27^−^ phenotype possessed suppressive activity. These results indicate that CD8⁺ Tregs belong to a narrower, more specialized regulatory subset rather than simply representing activated CD8⁺ T cells with NK-like traits. Accordingly, we propose that the phenotype of NKG2C^+^CD27^−^ distinguishes a distinct regulatory CD8⁺ T cell population from the heterogeneous population of NK-like CD8⁺ T cells, underscoring the need for refined subset definitions in studies of CD8⁺ regulatory immunity.

In this study, we were able to identify CD8⁺ Treg1 and Treg2 clusters within the CD45RA^+^CD27^−^ CD8^+^ Tsinglets obtained from the public CITE-seq and scTCR-seq dataset syn49637038 (Fig. 7A-C). We also attempted to identify CD8⁺ Treg clusters within total T cells and CD8⁺ T cells in this public dataset. However, this proved highly challenging or infeasible as CD8^+^ Tregs express a strong NK transcriptional program, encompassing *TYROBP*, *KLRC2*, *KIR* genes, etc. (Fig. 2C and D), which is also a feature of γδT cells (fig. S5B and S5C) (*70*). Furthermore, it is a common observation in scRNA-seq data analysis that a rare population—despite having a clean, specific cell surface marker profile by flow cytometry—may fail to form a distinct cluster in a UMAP projection. These two factors likely contributed to the observation that in a UMAP embedding of total T cells, Treg1 and Treg2 not only lacked distinct clusters but also partially overlapped with γδT cells (fig. S6D). Furthermore, even within a UMAP of extracted CD8^+^ T cells, Treg1 and Treg2 cells did not form separate clusters. Instead, they scattered into clusters of Tem, Tand NKT-like cells, due to shared activation/effector signature (fig. S6E). These results indicate that distinguishing CD8^+^ Tregs from these confounding populations requires protein-level CD45RA data to access the Tcompartment, and paired TCR sequencing to confirm αβ identity and exclude γδT cells, neither of which is available in most conventional scRNA-seq datasets. CD8^+^ Tregs will therefore be systematically misannotated in existing transcriptomic atlases, and their reliable identification requires either the NKG2C^+^CD27^−^ flow cytometry panel described here or multi-modal profiling integrating transcriptomics, surface proteomics, and TCR sequencing.

In summary, our study resolves prior heterogeneity in the field by establishing a unified, novel surface marker for CD8^+^ Tregs and their functionally distinct subsets, thereby providing a critical framework for their prospective isolation and in-depth functional investigation across health, aging, and disease.

## ACKNOWLEDGEMENTS

We thank Drs. Vijay K. Kuchroo, Jerome Ritz, Gordon J. Freeman, and Roni Nowarski for helpful discussion and suggestions. We thank Tian Cao and Alfred Laiman for processing discarded leukoreduction filters.

## Funding

1. D. H. was supported in part by National Multiple Sclerosis Society Research Grant RG-2111-38681 (to D.H.) and Brigham and Women’s Hospital Faculty Career Development Award (to D. H.). X.L. was supported in part by NIH/NIGMS T32GM007057-47 (to X.L.).

## Author contributions

Conceptualization: DH

Methodology: DH, XL, SKM

Data curation: SKM, DH, CT, RKK

Analysis and interpretation of data: DH, XL, SKM

Investigation: DH, XL, SKM

Visualization: XL, DH, SKM

Funding acquisition: DH, XL

Supervision: DH

Writing – original draft: DH, XL, SKM

Writing – review & editing: HLW, DH, XL, SKM, MH, RKK, TC

## Competing interests

A provisional patent application (US provisional application no. 64/046,734) covering the methods and data reported in this manuscript has been filed by Mass General Brigham with D.H., X.L., S.K.M. and H.W. listed as inventors. The authors/Institution may potentially benefit from future commercialization.

## Data, code, and materials availability

The authors declare that the main data supporting the findings of this study are available within the article and its Supplementary Information files. Additional data and code are available from the corresponding authors upon request. Single cell data will be deposited in a public database and released upon publication. The authors declare that there are no newly created materials in this study.

