## Supplemental Figures and Tables for "NKG2C^+^CD27^−^ Defines Human CD8^+^ Regulatory T Cells"

### SUPPLEMENTARY MATERIALS

### MATERTIALS AND METHODS

#### Human samples

Our studies received approval from the ethical standards committee on human studies at our institution (the Mass General Brigham Institutional Review Board). The four participants who provided blood samples for CITE-seq and scTCR-seq gave written informed consent. Peripheral blood was collected into sterile vacutainer tubes containing sodium heparin (BD Biosciences). Peripheral blood mononuclear cells (PBMCs) were isolated within 2 hours of collection by density gradient centrifugation using Ficoll® Paque Plus (Millipore Sigma) according to the manufacturer's instructions.

PBMCs used for flow cytometry and functional analyses were isolated from discarded leukoreduction filters obtained from Boston Children's Hospital. This work did not involve human subjects and therefore did not require IRB approval. Blood cells were recovered from used leukoreduction filters by flushing with Hank's Balanced Salt Solution (HBSS) supplemented with 2% heat-inactivated fetal bovine serum (FBS) and 2 mM EDTA. PBMCs were subsequently isolated by density gradient centrifugation using Ficoll® Paque Plus.

Isolated PBMCs were cryopreserved using freeze medium composed of 90% FBS and 10% dimethyl sulfoxide (DMSO) (sterile-filtered). Cells were adjusted to a final concentration of 1–5 × $10^7$ viable cells/mL in cryoprotectant solution. Aliquots of 1 mL were transferred into sterile, labeled cryovials (Corning). Cryopreserved cells were stored in a liquid nitrogen tank until use.

#### Flow cytometry

Cryopreserved PBMCs were thawed and cell viability was assessed by acridine orange/propidium iodide (AO/PI) staining. Only samples with ≥85% viability were used for downstream staining. Two custom antibody panels were developed to comprehensively identify and characterize human $CD8^+$ Tregs. Panel 1 (table S1), which included 13 antibodies, was designed for surface marker phenotyping to identify total $CD8^+$ T cells and $T_{EMRA}$, $CD56^+$ and $KIR^+$ populations based on established immunophenotypic signatures ($CD45RA^+CD27^-CCR7^-$, $CD56^+CD161^-$ and $KIR2DL2/L3/S2^+$ and/or $KIR3DL1^+$). Panel 2 (table S3), consisting of 24 antibodies, was developed based on Panel 1 and expanded to include novel cell surface markers (NKG2C, CD226, etc.) identified by our single-cell multi-omics analysis, enabling comprehensive characterization of $CD8^+$ Tregs. All antibodies were titrated prior to use to determine optimal concentrations.

For Panel 1, $1x10^6$ PBMCs per sample were stained with LIVE/DEAD™ Fixable Blue Stain (ThermoFisher Scientific, L23105), followed by incubating with Human TruStain FcX ™ (Fc Receptor Blocking Solution) (Biolegend, 422302). Antibody cocktail was prepared using BD Horizon™ Brilliant Stain Buffer Plus (WatersTM Biosciences, 566385) in the presence of True-Stain Monocyte Blocker ™ (Biolegend, 426102). Antibody cocktail was added to cells and incubated for 30 min at 4°C in the dark. Cells were washed twice with FACS buffer (DPBS containing 2% FBS). Stained cells were resuspended in FACS buffer and acquired on a Cytek Aurora cytometer equipped with 5 lasers (Cytek Biosciences). For each sample, 300,000 events were recorded. For Panel 2, $3x10^6$ PBMCs per sample were stained following a similar staining protocol, and 1,000,000 events were recorded per sample.

**$CD8^+$ Treg-mediated suppression assay**

Cryopreserved PBMCs were thawed and cell viability was assessed by AO/PI staining. Only samples with ≥90% viability were used. To prepare antigen-presenting cells (APCs), autologous and allogeneic PBMCs were treated with mitomycin C (MMC) (10 μg/ml) in a flat-bottom 96 well plate (1 x $10^6$ cells/ 50 μl/well) for 3 hours at 37°C in 5% $CO_2$ to prevent proliferation. After treatment, cells were washed three times with complete medium (IMDM medium supplemented with 10% FBS and 1% penicillin-streptomycin). MMC-treated autologous and allogeneic PBMCs were mixed at ratio of 1:1 and used as APCs in subsequent co-culture experiment.

Total T cells were isolated from (autologous) PBMCs using the EasySep™ Release Human CD3 Positive Selection Kit (StemCell Technologies, 17751). Isolated T cells were stained with fluorescence-conjugated antibodies for CD3, CD4, CD8, NKG2C, CD27, CD226, CD56, CD161, KIR2DL2/L3/S2, KIR3DL1, TCRγδ, and TCR Vα7.2 (table S4). Target pan T cells, total $CD8^+$ Tregs, $CD8^+$ Treg subsets and control $CD8^+$ T cells were sorted as described in fig. S5E. Target T cells were labeled with CellTrace Violet (5 μM) (Invitrogen, C34557).

Co-cultures were set up in 96-well U-bottom plates. The following cells were added per well, unless otherwise stated: APCs (1.6 x $10^4$), target pan T cells (8 x $10^3$) and $CD8^+$ Tregs or control T cells (2 x $10^3$). Co-cultures were established in nine experimental conditions: 1) APCs + Target cells + Control $CD8^+$ T cells, 2) APCs + Target cells + total $CD8^+$ Tregs (tTreg), 3) APCs + Target cells + Treg1, 4) APCs + Target cells + Treg2, 5) APCs + Target cells + $CD8^+$CD56+$CD161^-NKG2C^-$ T cells, 6) APCs + Target cells + $CD56^+NKG2C^+$ Tregs, 7) APCs + Target cells + $CD8^+$KIR+$NKG2C^-$ T cells, 8) APCs + Target cells + $KIR^+NKG2C^+$ Tregs, 9) APCs + Target cells only. All conditions were set up in triplicate wells. The final volume per well was adjusted to 200 μL with complete medium in the presence of IL-2 (50 unit/mL). Plates were incubated for 4 days at 37°C in 5% $CO_2$.

After the 4-day co-culture, cells were incubated with Human TruStain FcX for 10 minutes at room temperature, followed by staining with fluorescence-conjugated antibodies for CD3, CD4, CD8, NKG2C, CD27, CD226, CD56, CD161, KIR2DL2/L3/S2 and KIR3DL1. After staining, cells were washed twice with FACS buffer. Stained cells were resuspended in FACS buffer (175 μL /well) and acquired on a Cytek Aurora cytometer equipped with 5 lasers (Cytek Biosciences). For each sample, 120 μL /well of cell suspension were acquired.

**Cell sorting and library preparation for CITE-seq and V(D)J enrichment**

For single-cell analysis of $CD8^+$ T cells, we isolated $CD8^+$ $T_{EMRA}$ ($CD3^+CD8^+CD45RA^+CD27^-$) and $CD8^+$ $CD45RA^{neg}$ ($CD3^+CD8^+CD45RA^-$) cells from the peripheral blood mononuclear cells (PBMCs) of two healthy female (F) and two male (M) donors with a Cytek® Aurora CS cell sorter. An equal number of flow-sorted cells of each sample was pooled and multiplexed using hashtag antibodies: TotalSeq™-C0251 (CD8$T_{EMRA}$_F1), TotalSeq™-C0252 (CD8$T_{EMRA}$_F2), TotalSeq™-C0253 (CD8$T_{EMRA}$_M1), TotalSeq™-C0254 (CD8$T_{EMRA}$_M2), TotalSeq™-C0255 (CD8Teff_F1), TotalSeq™-C0256 (CD8Teff_F2), TotalSeq™-C0257 (CD8Teff_M1), and TotalSeq™-C0258 (CD8Teff_M2) (BioLegend) (table S2). The pooled cells were simultaneously stained with oligonucleotide-conjugated antibodies targeting CD4 (TotalSeq™-C0072), CD8α (TotalSeq™-C0080), CD45RA (TotalSeq™-C0063), CD45RO (TotalSeq™-C0087), CD56 (TotalSeq™-C0047), CD161 (TotalSeq™-C0149), CD16 (TotalSeq™-C0083), CD57 (TotalSeq™-C0168), CD183 (CXCR3) (TotalSeq™-C0140), and CD279 (PD-1) (TotalSeq™-C0088) (BioLegend) (table S2). Gene expression, surface protein expression, and TCR V(D)J repertoire profiling was performed using the Chromium GEM-X Single Cell 5' Reagent Kits v3 with Feature Barcode technology for cell surface protein (10x Genomics). Following a single cell capture on the 10x Chromium X controller, three separate libraries were prepared according to the manufacturer's instructions: gene expression (GEX), cell surface protein (CSP; including both antibody-derived tags [ADT] for proteins and hashtag oligos [HTO] for multiplexing), and TCR V(D)J. For V(D)J enrichment, the Human TCR Amplification Kit (16 rxns, PN-1000252) was used following the manufacturer's recommendations. All libraries were sequenced on an Illumina NovaSeq X Plus sequencer (PE150-25B, 1 lane).

**CITE-seq and V(D)J preprocessing with CellRanger**

Raw sequencing data were processed using CellRanger (version 8.0.1, 10x Genomics), mapping reads to the GRCh38 human reference genome (refdata-gex-GRCh38-2024-A). Gene expression (GEX) libraries were processed alongside antibody-derived tag (ADT) and hashtag oligonucleotide (HTO) feature barcoding libraries using the cellranger count pipeline with an expected cell recovery of ~40,000 cells per sample. T cell receptor (TCR) alpha and beta chain sequences were assembled using the cellranger vdj pipeline with the GRCh38 V(D)J reference (version 7.1.0). All libraries from the same sample were processed jointly to ensure barcode alignment across modalities. To prevent TCR transcripts from influencing downstream clustering, all TCR-specific genes were excluded from the filtered feature-barcode matrix using CellRanger reanalyze (v8.0.1), and a new count matrix was generated after removal of human TCR genes. V(D)J sequences were processed separately with CellRanger v8.0.1 using the human VDJ reference genome (refdata-cellranger-vdj-GRCh38-alts-ensembl-7.1.0). The CellRanger VDJ output (filtered_contig_annotations.csv) was then processed using custom code to identify the dominant TCR contig per cell. The TCR information per cell, including V(D)J gene segments, CDR3 sequences, and chain pairings, was merged with the gene expression (GEX) and cell surface protein (CSP) data as metadata columns in a Seurat object. This integration enabled multimodal downstream analysis, including correlating TCR clonotypes with gene expression profiles, protein marker levels, and sample origins for tasks such as clustering, differential expression, and clonal expansion assessment.

**Quality control, demultiplexing, integration and normalization of single-cell RNA-seq**

***Quality control and cell filtering***

Downstream analyses of CITE-seq were performed using Seurat (*71*) (version 5.2.0) in R (version 4.4.2). Gene expression data were log-normalized and cell cycle phase was assigned using the CellCycleScoring function with established S- and G2/M-phase gene markers. Quality control metrics were calculated for each cell, including mitochondrial gene content (%MT), ribosomal gene content (%RPL/RPS), and hemoglobin gene content (%HBA/HBB). Cells were retained if they had ≥900 unique genes detected, ≤75,000 total UMI counts, and ≤12% mitochondrial content.

***Sample demultiplexing***

HTO counts were normalized using centered log-ratio (CLR) transformation applied per cell. Sample identity was assigned using the HTODemux function in Seurat with a positive quantile threshold of 0.99. Cells classified as Doublet or Negative by HTODemux were excluded from downstream analyses.

***Data integration and normalization***

To account for subject-specific batch effects, data were integrated across subjects using Harmony (*72*) (max iterations = 20, $\theta = 4$, $\lambda = 0.5$, $\sigma = 0.2$). Initial clustering after integration identified contaminating B cells and monocytes, which were excluded based on expression of lineage-specific markers (e.g., CD19, CD14). $CD4^+$ T cells were identified and excluded based on anti-$CD4^+$ ADT signal (see $CD8^+$ T Cell Purification below). ADT data were normalized using CLR transformation applied per feature (margin = 2).

### Single-cell multi-omics analysis

#### *$CD8^+$ T cell extraction*

$CD8^+$ T cells were extracted in silico based on surface protein expression: $CD3^+CD4^-CD8^+$ as measured by ADT. To determine optimal expression cutoffs for CD4 and CD8, we fitted two-component Gaussian mixture models to the normalized ADT expression distribution of each marker using the mclust (*73*) package (version 6.1.2) in R. The intersection point between the two fitted Gaussian distributions was taken as the classification threshold for each marker.

#### *Dimensionality reduction and clustering*

$CD8^+$ T cells were normalized using SCTransform (v2), regressing out mitochondrial content percentage and cell cycle scores (S.Score and G2M.Score) as confounding variables. Principal component analysis (PCA) was performed on SCTransform-normalized data, and batch correction was applied to the top 30 principal components using Harmony with the same parameters described above. UMAP dimensionality reduction was computed from the first 30 Harmony-corrected principal components.

Cell clustering was performed using the Louvain community detection algorithm (resolution = 0.6) applied to a shared nearest neighbor (SNN) graph constructed from the Harmony-corrected PCA embedding. Cluster 7, which displayed characteristics of ambient RNA contamination with lower per-cell UMI counts relative to all other clusters (fig. S2C) was excluded from downstream analysis. MAIT cells were identified based on TCR gene usage (*TRAV1-2* paired with *TRAJ33/20/12*) confirmed by scTCR-seq data. All remaining clusters were annotated based on differential expression of established lineage and functional marker genes, protein-level ADT expression, and pathway enrichment analysis.

**Differential gene expression and pathway enrichment analysis**

Differentially expressed genes (DEGs) between $CD8^+$ T cell clusters were identified using the FindMarkers function in Seurat with a Wilcoxon rank-sum test. Genes were tested if expressed in ≥10% of cells in either comparison group (min.pct = 0.10). DEGs were defined by $|\log_2$ fold change$| > 0.25$ and Bonferroni-adjusted $p$-value < 0.05 unless otherwise specified. For the Treg2-versus-Treg1 comparison and the concordance analysis (Fig. 7C), a stricter threshold of $|\log_2 FC| > 0.50$ and FDR < 0.05 was applied.

For visualization, the top DEGs in the CITE-seq analysis of the isolated $CD8^+$ $T_{EMRA}$ and $CD8^+$ $CD45RA^{neg}$ cells were ranked by a composite score calculated as fold change multiplied by the percentage of positive cells (Fig. 2D), to prioritize markers that are both highly expressed and broadly detected within the cluster. Gene set enrichment and cell type annotation were performed using Enrichr (*46, 47*) (https://maayanlab.cloud/Enrichr) via the enrichR R package (version 0.3.0). Marker gene lists for each $CD8^+$ T cell cluster were submitted to Enrichr and queried against the CellMarker Augmented 2024 database for cell type identity annotation (Fig. 2C) and the Gene Ontology Biological Process 2023 database for pathway enrichment of DEGs between Treg2 and Treg1 (Fig. 3D). Results were filtered by adjusted $p$-value < 0.05 and ranked by combined score.

#### *Gene regulatory network inference*

Gene regulatory network (GRN) analysis was performed using pySCENIC (*74*) (version 0.12.1) following the standard three-step pipeline. First, co-expression modules (regulons) were inferred using arboreto_with_multiprocessing with GRNBoost2 and a curated list of 1,839 human transcription factors from the pySCENIC database. Second, co-expression modules were pruned based on cis-regulatory motif enrichment using the cisTarget database for the human genome (hg38), incorporating motif rankings for two region sets: (i) 10 kb upstream and downstream of transcription start sites (TSS), and (ii) 500 bp upstream to 100 bp downstream of the TSS. Motif annotations were drawn from the v10 motifs database, and technical dropout events were masked prior to scoring. Third, regulon activity in individual cells was quantified using AUCell with default parameters, generating a per-cell regulon activity score (AUC). Regulon specificity scores (RSS) were calculated for each annotated CD8+ T cell subset versus all remaining cells, and the top five regulons per subset ranked by RSS were selected for visualization.

#### *TCR repertoire analysis*

TCRα/β chain sequences were extracted from CellRanger VDJ filtered_contig_annotations.vsv output and processed using a custom R script. Briefly, V(D)J barcodes were first filtered to retain only those present in the corresponding gene expression (GEX) dataset. For each retained V(D)J barcode, productive contigs were separated by chain (TRA and TRB), and the dominant contig within each chain was defined as the contig with the highest read count. The corresponding TRA and TRB dominant contigs were used to assign the paired TCRα/β sequence annotations per cell. The resulting per-cell V(D)J annotations were then integrated into the preprocessed Seurat object (comprising GEX data with TCR genes removed, ADT, and HTO data) via barcode matching. A clonotype was defined as a unique combination of TCRα and TCRβ CDR3 amino acid sequences. Clonality was categorized by clone size as: singleton (n = 1), minimal (n = 2-4), moderate (n = 5-16), high (n = 17-64), and hyperexpanded (n > 64). TCR repertoire diversity was quantified using the Shannon entropy index, which accounts for both clonal richness (number of unique clonotypes) and evenness (distribution of clone sizes). Clonotype sharing across $CD8^+$ T cell subsets was visualized using UpSet plots. Pairwise repertoire overlap between subsets was quantified using the Morisita overlap index, which ranges from 0 (no shared clonotypes) to 1 (identical repertoires).

**Analysis of public single-cell multi-omics dataset**

To examine age-associated changes in $CD8^+$ Treg frequency, we reanalyzed a published CITE-seq and scTCR-seq dataset (accession: syn49637038) (*65*) comprising PBMCs from 166 healthy donors aged 25–85 years. Raw processed count matrices and ADT data were downloaded from Synapse and imported into Seurat for re-analysis.

Cell filtering and $CD8^+$ T cell enrichment were performed using ADT-based gating: γδ T cells (TCRγδ$^+$), MAIT cells (TCR Vα7.2$^+$), and $CD4^+$ T cells were excluded sequentially. High-quality $CD3^+CD4^-CD8^+$ singlets ($n$ = 271,875) were retained. $CD8^+$ $T_{EMRA}$ cells were further enriched by selecting $CD45RA^+CD27^-$ cells based on ADT expression ($n$ = 45,390), applying the same Gaussian mixture model approach described above to determine ADT expression cutoffs.

$CD8^+$ $T_{EMRA}$ cells were re-clustered using Harmony-corrected PCA followed by Louvain clustering (resolution = 0.6). Treg1 (Cluster 0) and Treg2 (Cluster 11) were identified based on co-expression of *TYROBP*, *KLRC2*, *IKZF2*, *KIR3DL1*, *KIR2DL3*, *KIR2DL1*, and *PTGDS* (for Treg2), consistent with the transcriptional signatures defined in our primary CITE-seq dataset.

Concordance of transcriptional signatures between datasets was assessed by independently calling DEGs ($|\log_2 FC| > 0.50$, FDR $< 0.05$) for the Treg2-versus- Treg1 comparison in each dataset. Directional concordance was evaluated by Pearson correlation of $\log_2$FC values across both datasets and visualized as a four-quadrant scatter plot.

The association between $CD8^+$ Treg subset frequency and donor age was assessed using linear regression. Cell frequencies among $CD8^+$ T cells or $CD8^+$ $T_{EMRA}$ cells were calculated per donor and plotted against age with Pearson correlation coefficients and *p*-values reported.

**Statistical Analysis**

All data are presented as mean ± s.d. unless otherwise noted. Statistical analyses of flow cytometric data were conducted using GraphPad Prism (version 10.5.0). The specific statistical test for each comparison is indicated in the corresponding figure legend. Paired, two-tailed Student's t-tests were used to compare paired samples from the same donor. Wilcoxon matched-pairs signed-rank tests were used for non-normally distributed paired data. Unpaired t-tests were used for between-group comparisons of independent samples. For comparisons across three or more groups, ordinary one-way ANOVA with Dunnett's multiple comparisons test (versus a designated control group) was applied. Pearson correlation was used to assess linear associations between continuous variables. A $p$-value $< 0.05$ was considered statistically significant throughout. For single-cell data, multiple testing correction was applied using the Bonferroni method (Seurat FindMarkers) or Benjamini–Hochberg FDR correction as specified per analysis.

### SUPPLEMENTARY FIGURES AND LEGEND

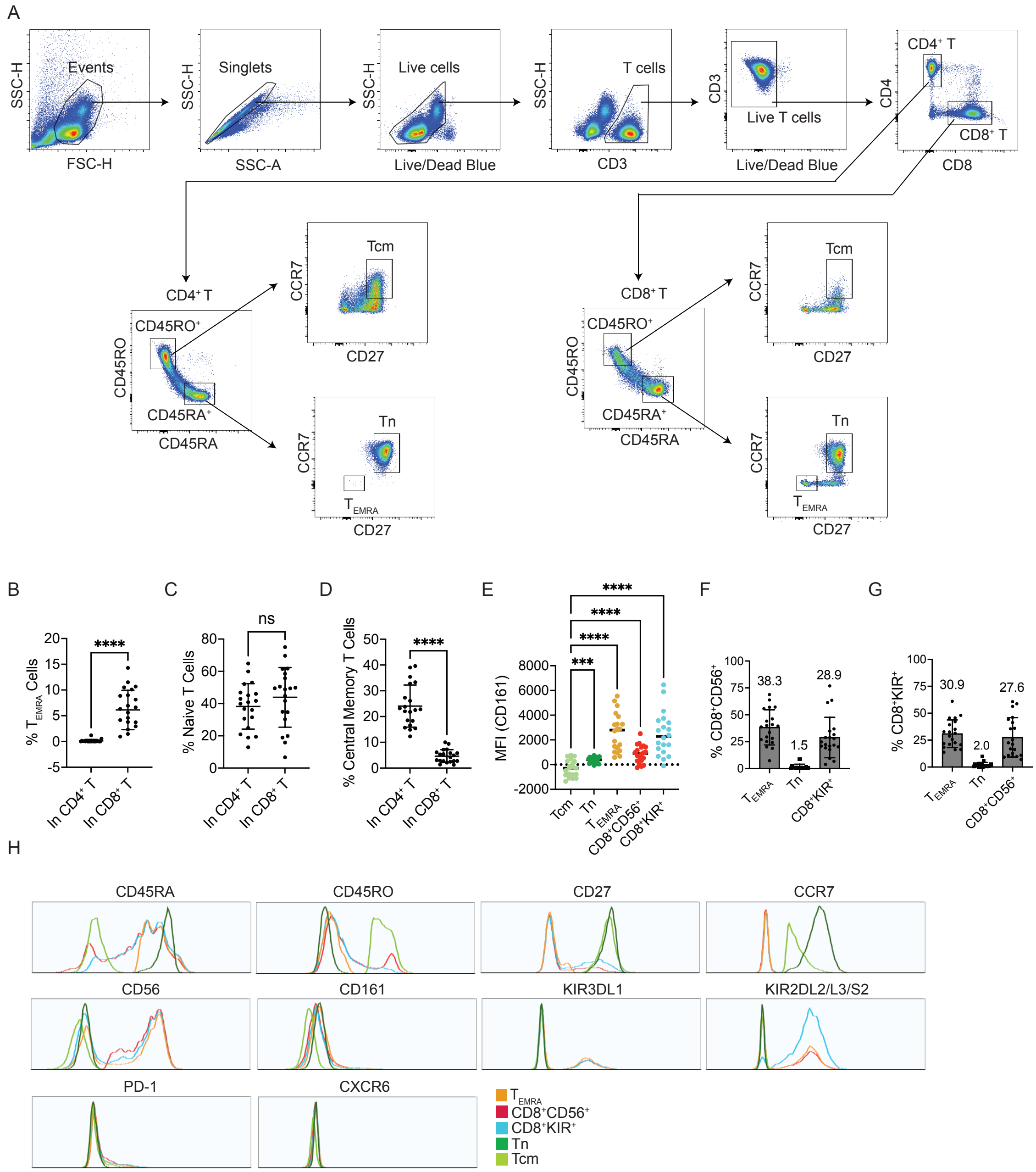


**fig. S1 Supplementary data on flow cytometry analysis for Figure 1.**

PBMCs from healthy donors (*n* = 20) were analyzed with flow cytometry. (**A**) Representative plots showing the gating strategy for CD4$^+$ and CD8$^+$ T cell subsets. (**B-D**) Plots for the frequency of T$_{EMRA}$ (**A**), Tn (**B**) and Tcm (**C**) cells in the CD4$^+$ and CD8$^+$ T cell compartments (mean ± s.d.).

Paired, two-tailed, *t*-test. ****, $p < 0.0001$; ns, not significant. (**E**) CD161 expression in $CD8^+$ T cell subsets. MFI, mean fluorescence intensity. Ordinary one-way ANOVA, Dunnett's multiple comparisons test, ***$p < 0.001$, ****$p < 0.0001$. (**F**) Frequencies of $T_{EMRA}$, Tn and $CD8^+KIR^+$ cells in the $CD8^+CD56^+$ compartments (mean ± s.d.). (**G**) Frequencies of $T_{EMRA}$, Tn and $CD8^+CD56^+$ cells in the $CD8^+KIR^+$ compartments (mean ± s.d.). (**H**) Representative histograms showing cell surface marker expression in CD8 T cell subsets. Tn, naïve T cell; Tcm, central memory T cell.

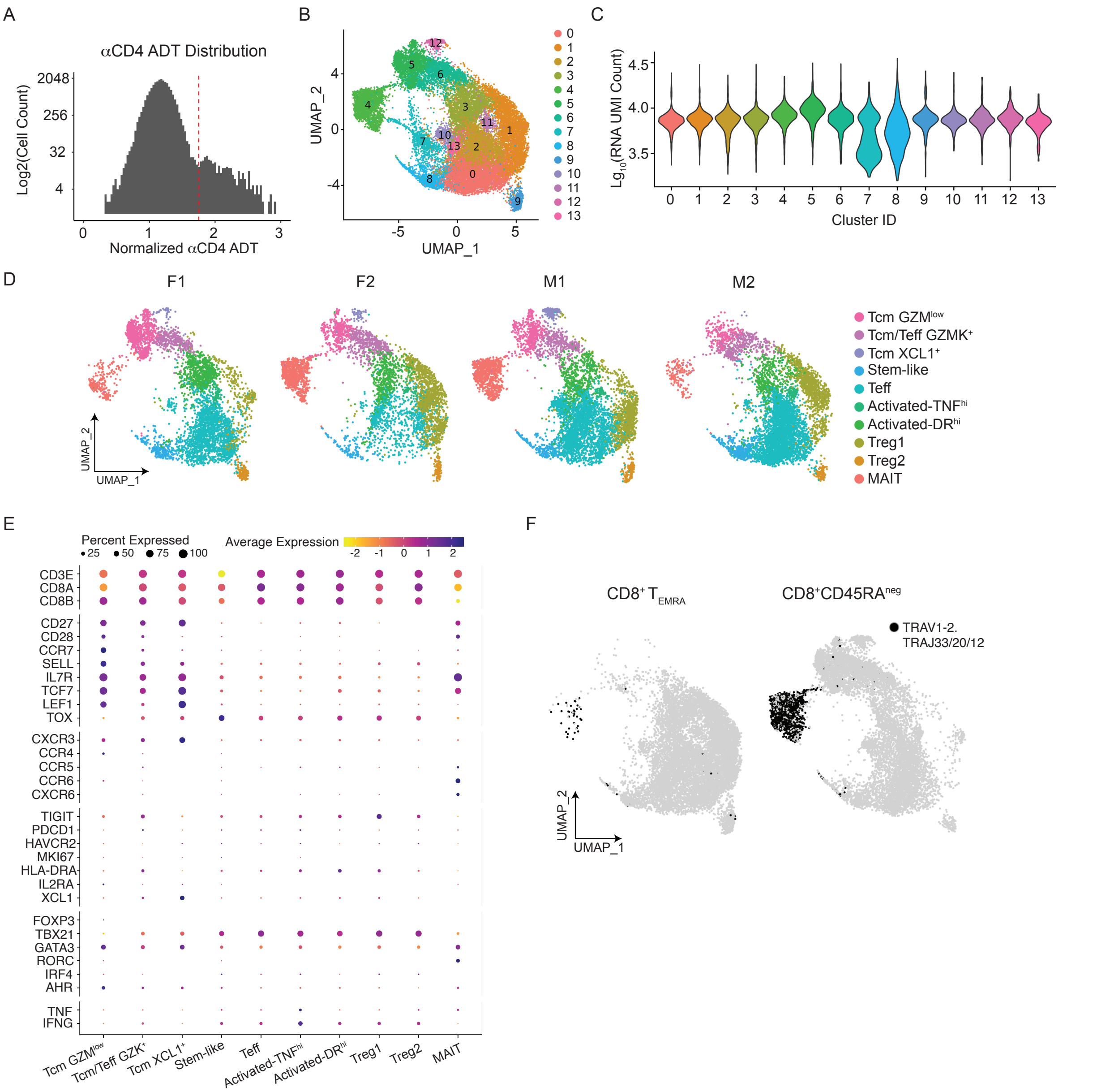


**fig. S2 Supplementary data on CITE-seq and scTCR-seq for Figure 1.**

Quality control (QC) filtering of CITE-seq and scTCR-seq data from CD8$^{+}$ T$_{EMRA}$ and CD45RA$^{neg}$ cells isolated from PBMCs of HCs (*n* = 4; 2 females, 2 males) to retain high-quality singlets, with per-sample UMAPs confirming successful integration across samples. (**A**) Histogram showing the distribution of cells positive for anti-CD4-ADT, which were excluded for downstream analysis.

(**B**) UMAP plot of $CD8^+$ T cells after excluding $CD4^+$ T cells, colored by the 14 identified clusters. (**C**) Violin plot of RNA UMI counts per cluster revealed that Cluster 7 had substantially lower counts. Cells from this cluster were therefore excluded from downstream analysis due to low quality. (**D**) UMAP plots of high quality $CD8^+$ T cell singlets from each donor. F, female; M, male.

(**E**) Expression of $CD8^+$ T cell subset-related genes across $CD8^+$ T cell clusters. Dot color indicates the level of gene expression and dot size indicates the proportion of cells expressing the gene. (**F**) UMAP plots illustrating the distribution of MAIT cells within $CD8^+$ $T_{EMRA}$ ($n$ = 11,181) and $CD45RA^{neg}$ ($n$ = 11,490) cells, respectively. Each black dot represents a *TRAV1-2*$^+$*TRAJ33/20/12*$^+$ MAIT cell.

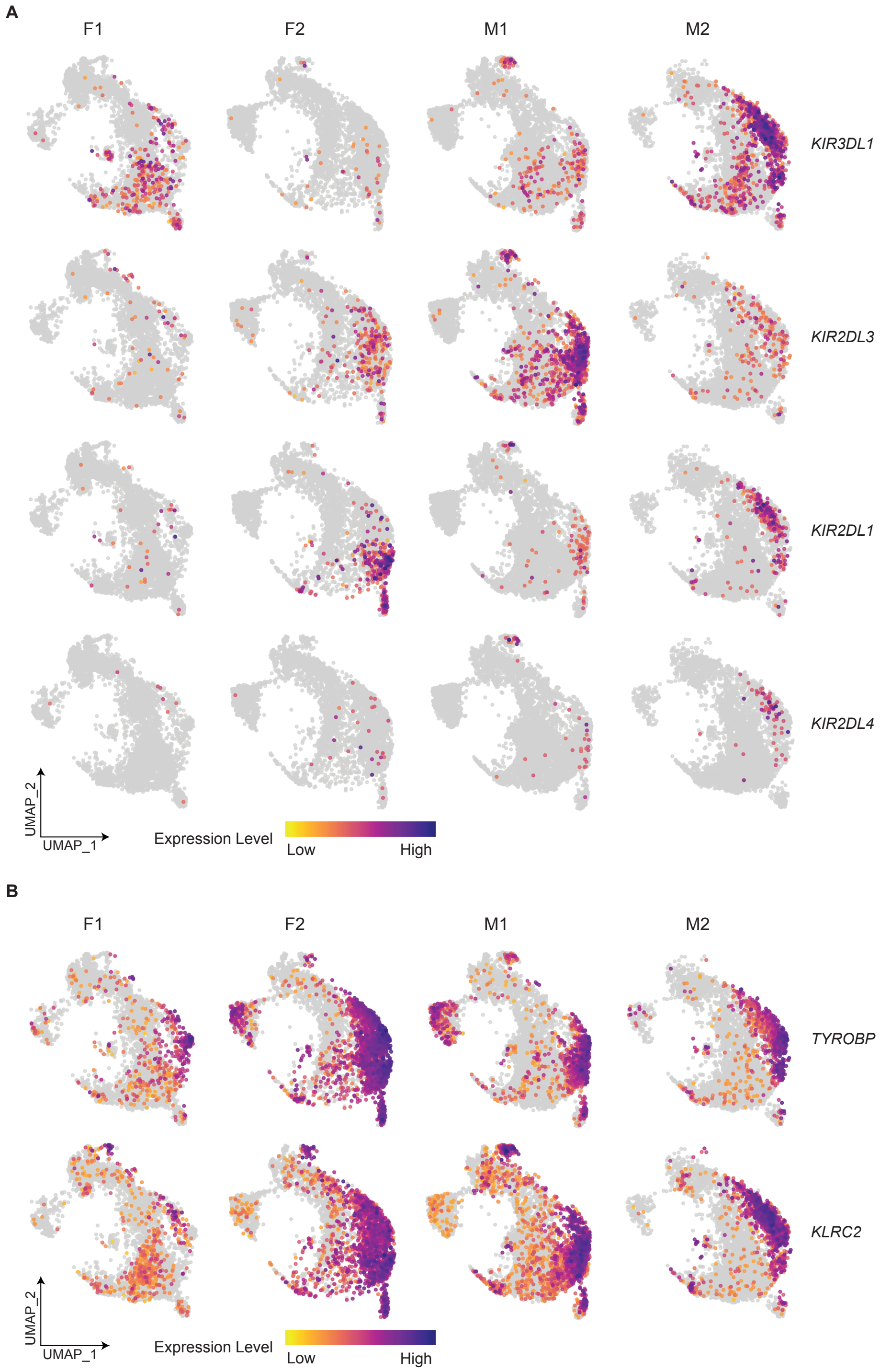


**fig. S3 Supplementary data on CITE-seq for Figure 2.**

(**A**) UMAP plots of individual donors showing stochastic expression of representative KIR genes across donors. (**B**) UMAP plots showing the expression of *TYROBP* and *KLRC2* across donors.

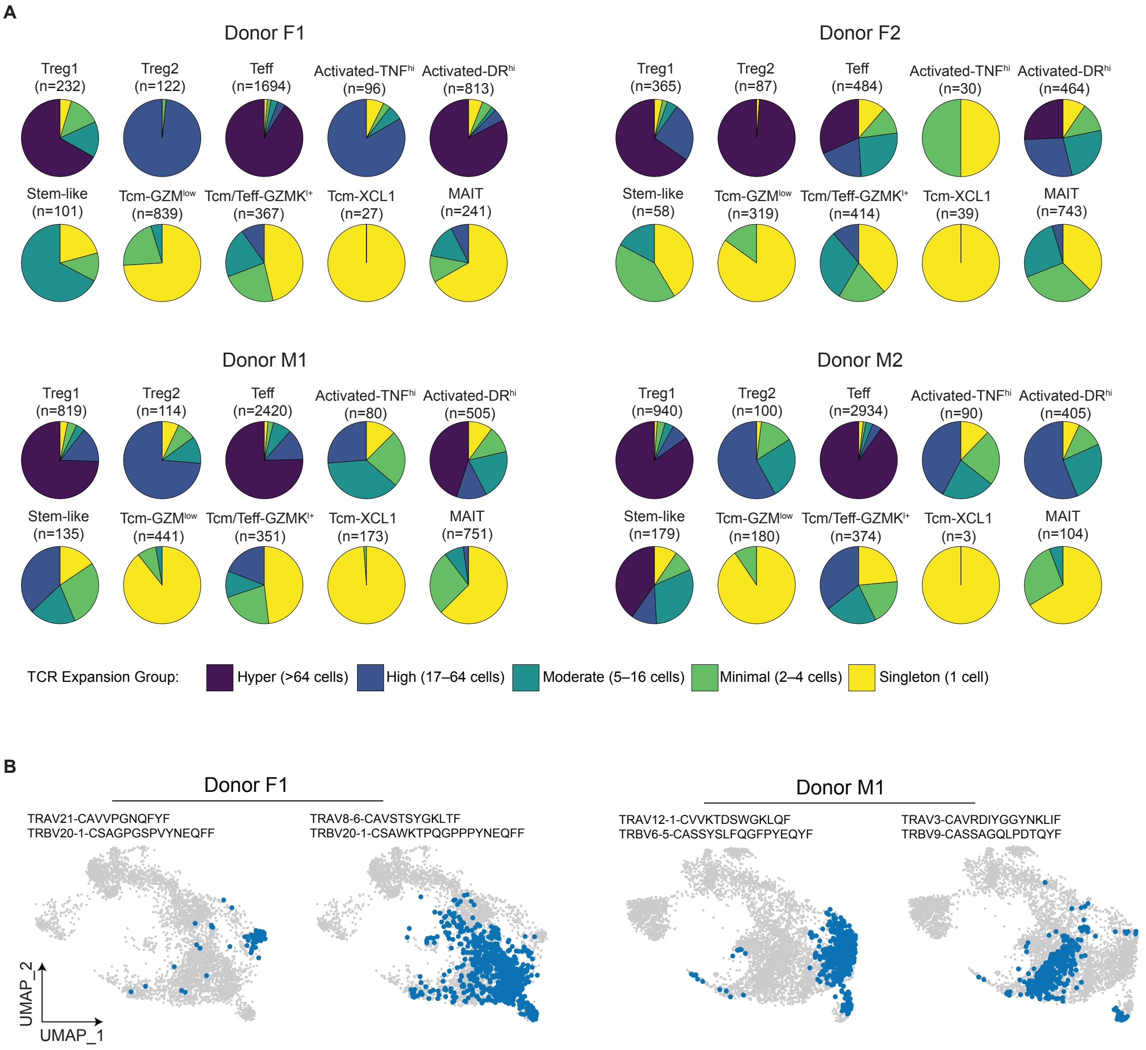


**fig. S4 Supplementary data on scTCR-seq for Figure 4.**

(**A**) Relative abundance of cells with varying degrees of clonality (singleton, minimal, moderate, high and hyper) within each CD8$^{+}$ T cell subpopulation across donors. The number above each pie chart indicates the number of cells in the corresponding subpopulation. (**B**) UMAP plots of merged CD8$^{+}$ T$_{EMRA}$ and CD45RA$^{neg}$ cells of donor F1 (left panel) and donor M1 (right panel). Four representative TCRαβ clonotypes are depicted, each of which shared between Treg1 and/or Treg2 cells and other $CD8^+$ T cell subpopulations. Blue dots represent individual cells expressing the TCRαβ clonotype indicated above the corresponding UMAP plot.

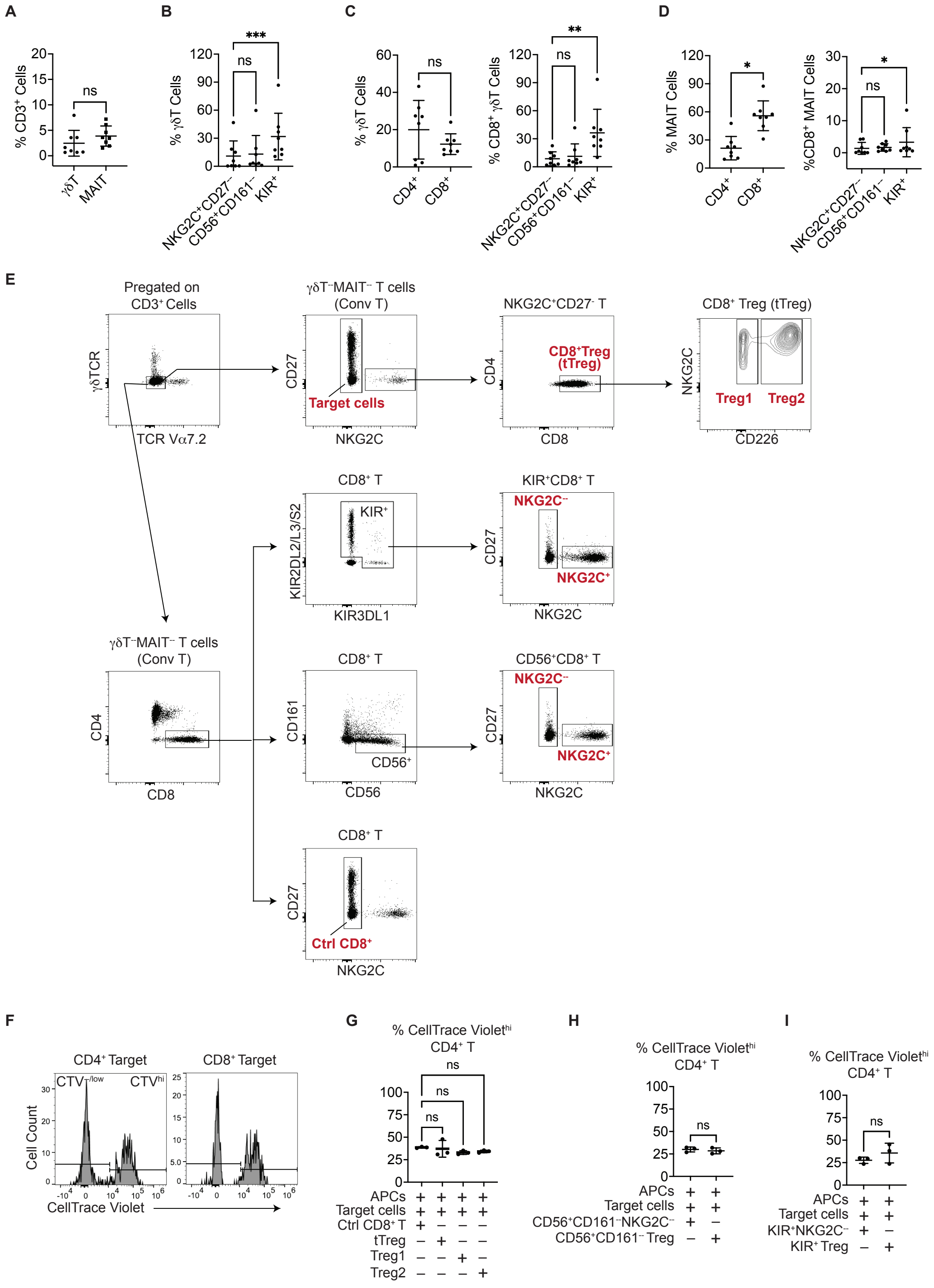


**fig. S5 Supplementary data for Figure 5 and 6.**

(**A**-**D**) Supplementary data for Figure 5. PBMCs from HCs were analyzed using a CyTek® Aurora flow cytometer with a 24-antibody panel ($n$ = 8, mean ± s.d.). (**A**) Frequencies of γδT cells and MAIT cells within the $CD3^+$ T cell compartment. Wilcoxon matched-pairs signed-rank test. (**B**) Frequencies of $NKG2C^+CD27^–$, $CD56^+CD161^–$ and $KIR^+$ cells within the γδT cell compartment. Ordinary one-way ANOVA, Dunnett's multiple comparisons test, *** $p < 0.001$. (**C**) Left, frequencies of $CD4^+$ and $CD8^+$ T cells within the γδT cell compartment. Wilcoxon matched-pairs signed-rank test. Right, frequencies of $NKG2C^+CD27^–$, $CD56^+CD161^–$ and $KIR^+$ cells within the γδT cell compartment. Ordinary one-way ANOVA, Dunnett's multiple comparisons test, ** $p < 0.01$. (**D**) Left, frequencies of $CD4^+$ and $CD8^+$ T cells within the MAIT cell compartment. Wilcoxon matched-pairs signed-rank test. Right, frequencies of $NKG2C^+CD27^–$, $CD56^+CD161^–$ and $KIR^+$ cells within the MAIT cell compartment. Ordinary one-way ANOVA, Dunnett's multiple comparisons test, * $p < 0.05$. (**E**-**H**) Supplementary data of the $CD8^+$ Treg-mediated suppression assay for Figure 6. (**E**) Sorting scheme of isolating cells for $CD8^+$ Treg-mediated suppression assay. Cell subsets marked in red were collected for the assay. (**F**) Histograms of pregated $CD4^+$ and $CD8^+$ T cells of CTV-labeled $CD3^+$ target cells co-cultured with APCs in the absence of $CD8^+$ Tregs for 4 days, showing cell proliferation indicated by the $CTV^{-/low}$ proliferation. In **G**-**I**, cells co-cultured in the presence of $CD8^+$ Tregs were harvested on day 4 and analyzed by flow cytometry, and the x-axis indicates the co-culture conditions ($n$ = 3, mean ± s.d.). (**G**) Frequencies of CellTrace $Violet^{hi}$ cells among $CD4^+$ T cells after co-culture with control $CD8^+NKG2C^–$, tTreg, Treg1 or Treg2 cells. Ordinary one-way ANOVA, Dunnett's multiple comparisons test. (**H**) Frequencies of CellTrace $Violet^{hi}$ cells among $CD4^+$ T cells after co-culture with control $CD8^+NKG2C^–CD56^+CD161^–$ or $CD56^+NKG2C^+$ Treg cells. Unpaired $t$-test. (**I**)

Frequencies of CellTrace Violet$^{hi}$ cells among $CD4^+$ T cells after co-culture with control $CD8^+NKG2C^-KIR^+$ or $KIR^+NKG2C^+$ Treg cells. Unpaired *t*-test.

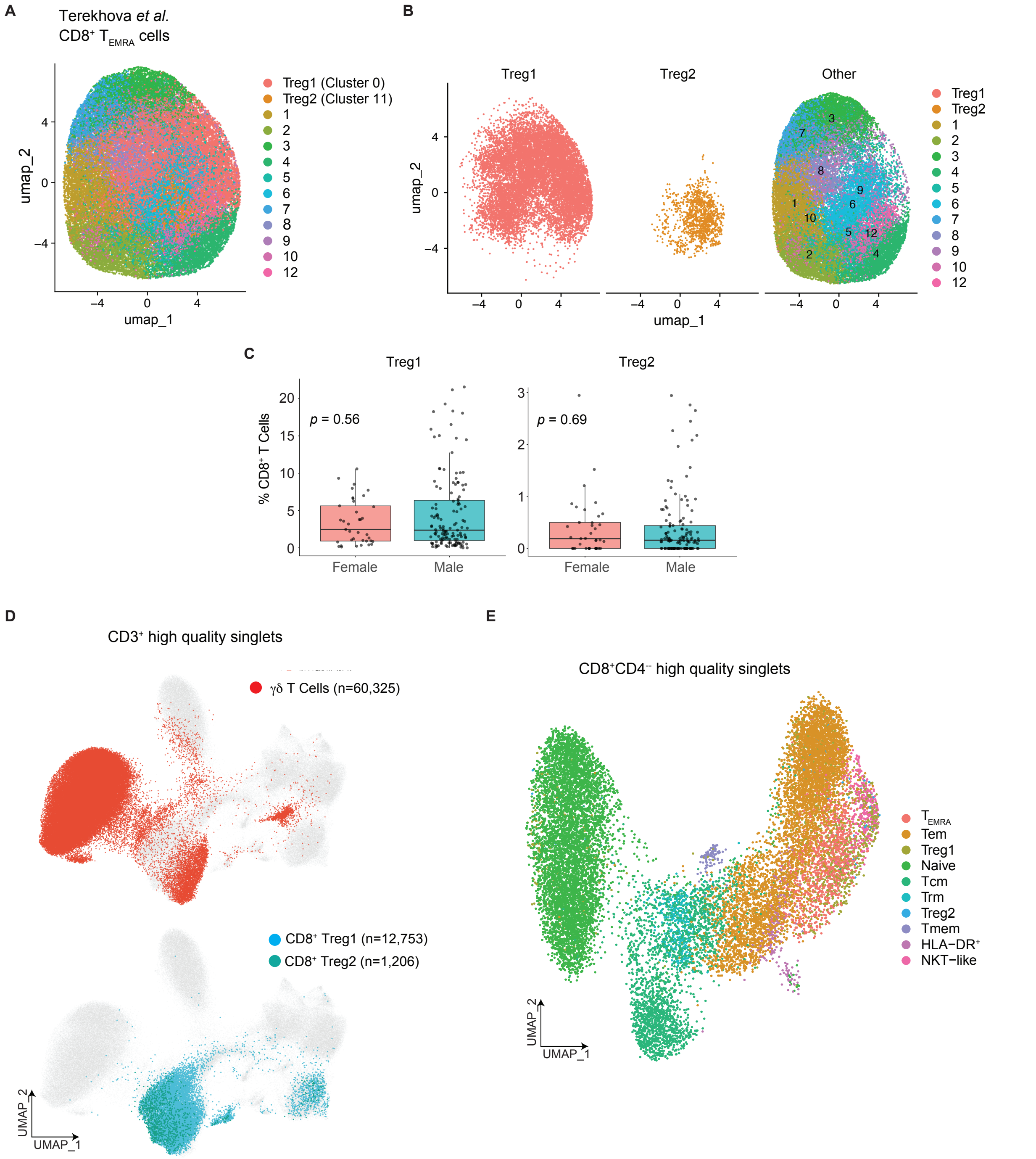


**fig. S6 Supplementary data for Figure 7.**

Data source: public scRNA-seq dataset syn49637038. (**A**) UMAP plot of CD8$^+$ T$_{EMRA}$ cells, colored by the 13 identified clusters, with Cluster 0 annotated as Treg1 and Cluster 11 as Treg2.

(**B**) Split UMAPs showing the distribution of Treg1 (left, 12,753 cells), Treg2 (middle, 1,206 cells) and the rest of $CD8^+$ $T_{EMRA}$ cells (right). (**C**) No sex-dependent differences in the frequencies of Treg1 and Treg2 were observed. Unpaired *t*-test ($n$ = 166). (**D**) UMAPs showing the distribution of γδT cells (upper) and $CD8^+$ Treg1 and Treg2 cells (lower) among $CD3^+$ T cells. (**E**) UMAP showing the distribution of Treg1 and Treg2 cells among $CD8^+$ T cells.

### SUPPLEMENTRAY TABLES AND LEGEND

### table S1 Antibody Panel 1 for Flow Cytometry Analysis.

| Antigen | Clone | Fluors |
|---|---|---|
| CD3 | UCHT1 | Brilliant Violet 510 |
| CD4 | RPA-T4 | Brilliant Violet 605 |
| CD8 | 3B5 | Qdot 800 |
| CD45RA | HI100 | Alexa Fluor 700 |
| CD45RO | UCHL1 | Brilliant Violet 711 |
| CD56 | NCAM16.2 | BUV737 |
| CD161 | HP-3G10 | APC/Fire 750 |
| CD197/CCR7 | G043H7 | Brilliant Violet 421 |
| CD158b/j(KIR2DL2/L3/S2) | DX27 | APC |
| CD158e1(KIR3DL1) | DX9 | PE |
| CD27 | M-T271 | Pacific Blue |
| CD279 (PD-1) | A17188A | FITC |
| CXCR6 (CD186) | K041E5 | PerCP/Cy5.5 |

### table S2 TotalSeq Antibodies for CITE-seq.

| Hashing Antibodies | Clone | Barcode |
|---|---|---|
| TotalSeq™-C0251 anti-human Hashtag 1 | LNH-94, 2M2 | GTCAACTCTTTAGCG |
| TotalSeq™-C0252 anti-human Hashtag 2 | LNH-94, 2M2 | TGATGGCCTATTGGG |
| TotalSeq™-C0253 anti-human Hashtag 3 | LNH-94, 2M2 | TTCCGCCTCTCTTTG |
| TotalSeq™-C0254 anti-human Hashtag 4 | LNH-94, 2M2 | AGTAAGTTCAGCGTA |
| TotalSeq™-C0255 anti-human Hashtag 5 | LNH-94, 2M2 | AAGTATCGTTTCGCA |
| TotalSeq™-C0256 anti-human Hashtag 6 | LNH-94, 2M2 | GGTTGCCAGATGTCA |
| TotalSeq™-C0257 anti-human Hashtag 7 | LNH-94, 2M2 | TGTCTTTCCTGCCAG |
| TotalSeq™-C0258 anti-human Hashtag 8 | LNH-94, 2M2 | CTCCTCTGCAATTAC |
| **CITE-seq Antibodies** | | |
| TotalSeq™-C0072 anti-human CD4 | RPA-T4 | TGTTCCCGCTCAACT |
| TotalSeq™-C0080 anti-human CD8α | RPA-T8 | GCTGCGCTTTCCATT |
| TotalSeq™-C0063 anti-human CD45RA | HI100 | TCAATCCTTCCGCTT |
| TotalSeq™-C0087 anti-human CD45RO | UCHL1 | CTCCGAATCATGTTG |
| TotalSeq™-C0047 anti-human CD56 | 5.1H11 | TCCTTTCCTGATAGG |
| TotalSeq™-C0149 anti-human CD161 | HP-3G10 | GTACGCAGTCCTTCT |
| TotalSeq™-C0083 anti-human CD16 | 3G8 | AAGTTCACTCTTTGC |
| TotalSeq™-C0168 anti-human CD57 | QA17A04 | AACTCCCTATGGAGG |
| TotalSeq™-C0140 anti-human CXCR3 | G025H7 | GCGATGGTAGATTAT |
| TotalSeq™-C0088 anti-human PD-1 | EH12.2H7 | ACAGCGCCGTATTTA |

**table S3 Antibody Panel 2 for Flow Cytometry Analysis.**

| Antigen | Clone | Fluors |
|---|---|---|
| CD3 | UCHT1 | Brilliant Violet 510 |
| CD4 | RPA-T4 | PerCP |
| CD8 | 3B5 | Qdot 800 |
| CD45RA | HI100 | BUV395 |
| CD45RO | UCHL1 | Brilliant Violet 711 |
| CD27 | L128 | BB700 |
| CD62L | DREG-56 | Brilliant Violet 650 |
| CD122 | TU27 | PE-Dazzle 594 |
| CD28 | CD28.2 | Brilliant Violet 605 |
| CD56 | NCAM16.2 | BUV737 |
| CD161 | DX12 | R718 |
| CD158b/j(KIR2DL2/L3/S2) | DX27 | PE |
| CD158e1(KIR3DL1) | DX9 | Brilliant Violet 421 |
| NKG2C | S19005E | FITC |
| NKG2A | S19004C | PE/Cy5 |
| NKp80 | 5D12 | APC |
| NKp46 | 9E2 | Alexa Fluor® 647 |
| TCR Vα7.2 | 3C10 | Spark Blue 574 |
| TCRγδ | B1.1 | PerCP-eF710 |
| PD-1 | A17188B | PE/Fire 810 |
| Tim-3 | F38-2E2 | Pacific Blue |
| CD226 | 11A8 | APC/Fire 750 |
| CD160 | BY55 | PE/Cyanine7 |
| CD197/CCR7 | G043H7 | Brilliant Violet 785 |

**table S4 Antibody Panel 3 for CD8$^+$ Treg Sorting.**

| Antigen | Clone | Fluors |
|---|---|---|
| CD3 | UCHT1 | Brilliant Violet 510 |
| CD4 | RPA-T4 | PerCP |
| CD8 | 3B5 | Qdot 800 |
| CD27 | L128 | BB700 |
| CD56 | NCAM16.2 | BUV737 |
| CD161 | DX12 | R718 |
| CD158b/j(KIR2DL2/L3/S2) | DX27 | APC |
| CD158e1(KIR3DL1) | DX9 | Brilliant Violet 421 |
| NKG2C | S19005E | FITC |
| TCR Vα7.2 | 3C10 | Spark Blue 574 |
| TCRγδ | B1.1 | PerCP-eF710 |
| CD226 | 11A8 | PE |

**List of references only cited in the Supplementary Materials**


71. Y. Hao *et al.*, Dictionary learning for integrative, multimodal and scalable single-cell analysis. *Nat Biotechnol* **42**, 293-304 (2024).
72. I. Korsunsky *et al.*, Fast, sensitive and accurate integration of single-cell data with Harmony. *Nat Methods* **16**, 1289-1296 (2019).
73. L. Scrucca, M. Fop, T. B. Murphy, A. E. Raftery, mclust 5: Clustering, Classification and Density Estimation Using Gaussian Finite Mixture Models. *R J* **8**, 289-317 (2016).
74. B. Van de Sande *et al.*, A scalable SCENIC workflow for single-cell gene regulatory network analysis. *Nat Protoc* **15**, 2247-2276 (2020).

**List of figures**

Fig. 1 $CD8^+$ Tregs are highly enriched within the $T_{EMRA}$ cells population and can be divided into two distinct subsets.

Fig. 2 Identification of TYROBP and NKG2C (*KLRC2*) as novel and improved markers for $CD8^+$ Tregs.

Fig. 3 *PTGDS* (encodes prostaglandin D2 synthase) and *CD226*: key markers distinguishing $CD8^+$ Treg2 from $CD8^+$ Treg1.

Fig. 4 $CD8^+$ Tregs exhibit high clonality and share T-cell receptors with diverse $CD8^+$ T cell subpopulations.

Fig. 5 Flow cytometric validation: $NKG2C^+CD27^-$ defines a $CD8^+$ Treg-specific phenotype, while CD226 differentiates Treg1 and Treg2.

Fig. 6 $CD8^+$ Tregs suppress $CD8^+$ T cells *in vitro*, with Treg2 showing stronger suppressive activity than Treg1.

Fig. 7 The frequency of $CD8^+$ Treg2 increases with aging among $CD8^+$ $T_{EMRA}$ cells.
